# Imaging genetics of language network functional connectivity reveals links with language-related abilities, dyslexia and handedness

**DOI:** 10.1101/2023.11.22.568256

**Authors:** Jitse S. Amelink, Merel C. Postema, Xiang-Zhen Kong, Dick Schijven, Amaia Carrion Castillo, Sourena Soheili-Nezhad, Zhiqiang Sha, Barbara Molz, Marc Joliot, Simon E. Fisher, Clyde Francks

**Affiliations:** Language and Genetics Department, Max Planck Institute for Psycholinguistics, Nijmegen, The Netherlands; Department of Psychology and Behavioural Sciences, Zhejiang University, Hangzhou, China; Department of Psychiatry of Sir Run Shaw Hospital, Zhejiang University School of Medicine, Hangzhou, China; Groupe d’Imagerie Neurofonctionnelle, Institut des Maladies Neurodégénératives, UMR5293, Commissariat à l’énergie atomique et aux énergies alternatives, CNRS, Université de Bordeaux, Bordeaux, France; Donders Institute for Brain, Cognition and Behaviour, Radboud University, Nijmegen, The Netherlands; Department of Cognitive Neuroscience, Radboud University Medical Center, Nijmegen, The Netherlands

## Abstract

Language is supported by a distributed network of brain regions with a particular contribution from the left hemisphere. A multi-level understanding of this network requires studying its genetic architecture. We used resting-state imaging data from 29,681 participants (UK Biobank) to measure connectivity between 18 left-hemisphere regions involved in multimodal sentence-level processing, as well as their right-hemisphere homotopes, and interhemispheric connections. Multivariate genome-wide association analysis of this total network, based on genetic variants with population frequencies *>* 1%, identified 14 genomic loci, of which three were also associated with asymmetry of intrahemispheric connectivity. Polygenic dispositions to lower language-related abilities, dyslexia and left-handedness were associated with generally reduced leftward asymmetry of functional connectivity. Exome-wide association analysis based on rare, protein-altering variants (frequencies *≤* 1%) suggested 7 additional genes. These findings shed new light on genetic contributions to language network organization and related behavioural traits.

## Introduction

The degree of sophistication in verbal communicative capacities is a uniquely defining trait of human beings compared to other primates. A distinctive feature of the neurobiology of language is its hemispheric dominance. Language lateralization starts when language is learned and intensifies during development into adulthood [1], resulting in leftward hemispheric dominance in about 85 percent of adults [2]. Most remaining adults have no clear dominant hemisphere for language, while roughly one percent show rightward hemispheric language dominance [2]. The left-hemisphere language network comprises various distributed regions including hubs in the inferior frontal gyrus and superior temporal sulcus[3, 4]. However, to a lesser extent, the right hemisphere homotopic regions are also active during language tasks, especially during language comprehension rather than production[3].

Language-related cognitive performance is highly heritable [5–10], and genetic factors also play a substantial role in susceptibility to language-related neurodevelopmental disorders such as childhood apraxia of speech [11], developmental language disorder (previously referred to as specific language impairment) and dyslexia [12–14]. In addition, hemispheric dominance for language builds on structural and functional asymmetries that are already present in neonates [15]. This suggests an early developmental basis for such asymmetries that is driven by a genetic developmental program [16–18].

Genome-wide association studies (GWAS) in tens or hundreds of thousands of individuals have begun to identify individual genomic loci associated with language- and/or reading-related performance [9], dyslexia [14], brain structural asymmetry [18] and/or left- or mixed-handedness [19]. Handedness is a behavioual manifestation of brain asymmetry with subtle and complex relations to hemispheric language dominance and language-related cognition and disorders [2, 14, 20]. The implicated genes in these GWAS tend to be most strongly expressed in the embryonic and fetal brain rather than postnatally. All together, these findings suggest that genetic contributions to inter-individual variation in language-related performance, and functional and structural brain asymmetries, exert their effects mostly early in life.

The genetic variants identified so far explain only a small proportion of the heritable variance in language-related performance or its structural underpinnings in the brain. A complementary approach to finding genes involved in language is to measure functional connectivity within the network of regions that support language in the brain, in many thousands of individuals, in order to perform well-powered GWAS. There are no existing datasets of this size that have collected functional imaging data during language task performance, but resting state functional connectivity is predictive of task-related functional activation [21–23] and also reveals meaningful organisation of the human cortex [24, 25]. The resting state functional connectivity approach involves identifying similarities between different brain regions in terms of their time course variation in the deoxyhemoglobin to hemoglobin ratio during the resting state, i.e. while participants are awake but not performing any particular task during functional magnetic resonance imaging (fMRI). The task-free nature of resting state fMRI makes it insensitive to choices in task design that can affect lateralization estimates [26], and is potentially more useful for studying the language network as a whole rather than circuits activated by one specific task. In addition, task-based fMRI has tended to find generally less heritable measures compared to resting state fMRI [27], making the latter perhaps more suitable for genetic investigation.

Previous work by Mekki et al. [28] found 20 loci in a genome-wide association study of functional language network connectivity based on resting state fMRI. The 25 brain regions used in their analyses to capture the brain’s language network were defined based on a meta-analysis of language-task activation across multiple previous task fMRI studies [29]. Of these 25 brain regions, 20 are in the left hemisphere and only 5 in the right hemisphere. The 25 regions were then analysed jointly with no further attention to hemispheric differences. However, given the early developmental basis of functional asymmetries [15], we reasoned that it may be informative for genetic association analysis to consider connectivity and hemispheric differences between all bilateral pairs of involved regions. For the present study we therefore chose a functional atlas with left and right hemisphere homotopies [30], developed in the BIL&GIN cohort, which consists of approximately 300 young adults roughly balanced for handedness. In previous work in this cohort, a core language network was defined in right handers (N=144) based on three language tasks (reading, listening, and language production) and a resting state paradigm [3]. A consensus multimodal language network called SENSAAS was defined, consisting of 18 regions in the left hemisphere that were active during all three language tasks.

For the purpose of the present gene mapping study, the right hemisphere homotopic regions were also included, yielding 36 regions in total (18 per hemisphere). We derived functional connectivity measures between these 36 regions (figure S1) in 29,681 participants from the UK Biobank who had genetic and brain imaging data available, yielding 630 intra- and interhemispheric connectivity measures and 153 hemispheric differences between left and right intrahemispheric connectivity. We then investigated multivariate associations of these functional connectivity phenotypes with common genetic variants, as well as polygenic scores for language-related abilities [9], dyslexia [14] and left-handedness [19].

In addition, we hypothesized that rare, protein-altering variants could also contribute to functional language connectivity, with relatively large effects in the few people who carry them. Such variants could give more direct clues to biological mechanisms underlying the formation of the brain’s language network. Previous large-scale genetic studies of both brain [20, 28] and cognitive or behavioural language-related traits [9, 10, 14] only analyzed common genetic variants (allele frequency in the population *≤* 1%). Tentative evidence for rare variant associations with right-hemisphere language dominance, involving actin cytoskeleton genes, was found in an exploratory study of 66 unrelated participants [31]. The first exome-wide association studies of the UK Biobank [32, 33] included structural brain imaging metrics, but not functional metrics. Therefore, the possible contributions of rare protein-coding variants to functional language connectivity had yet to be investigated in a biobank-sized data set, prior to the present study. After quality control (see Methods) we included 29,681 participants from the UK Biobank between ages 45 and 82 years, for whom single nucleotide polymorphism (SNP) genotyping array data, exome sequences, and resting state fMRI data were available, and that were in a previously defined ’white British’ ancestry cluster [34] (by far the largest single cluster in the data set). For these participants we derived 630 Pearson correlations between the time courses of the 36 regions in the language network (hereafter language network edges) and 153 hemispheric differences between left and right intrahemispheric homotopies (L-R, hereafter hemispheric differences) (figure S1). Positive hemispheric differences correspond to stronger connectivity on the left and negative hemispheric differences correspond to stronger connectivity on the right. We excluded language nework edges or hemispheric differences with no significant heritability (nominal p *≤* 0.05) (see figure S2 and Methods section), which left 629 edges and 103 hemispheric differences (table S1), among which the median SNP-based heritability was 0.070 (min: 0.018, max: 0.165) for language network connectivity and 0.026 (min: 0, max: 0.070) for hemispheric differences.

### Common genetic variant associations with language network connectivity and asymmetry

The 629 language network edges were entered into a multivariate genome-wide association scan (mvGWAS) with 8,735,699 biallelic SNPs (genome build hg19) that passed variant quality control (see Methods), using the MOSTest software [35] (see Methods), after controlling for potential confounders including age and sex (Methods). Using the standard GWAS multiple comparison threshold (5 *×* 10*^−^*^8^), 14 independent genomic loci showed significant multivariate associations with language network edges (figure 1A, table S2, figure S3). Subsequent gene mapping based on positional, eQTL and chromatin interaction information of SNPs (using FUMA [36]) found 111 associated genes (of which 40 were protein-coding, table S3). In addition, tissue expression analysis with MAGMA [37] showed preferential expression of language network associated genetic effects in prenatal development in the Brainspan gene expression data [38], which was significant at 21 weeks post conception but also generally elevated prenatally (figure 1C). Enrichment analysis against 11,404 gene sets (gene ontology and other curated sets) [39, 40] found no significant associations after correction for multiple comparisons, and cross-tissue enrichment analysis with respect to postmortem whole-body expression levels from GTEx [41] also found no significantly higher expression in any particular tissue of the body (figure S4 and table S4).

**Figure 1:**
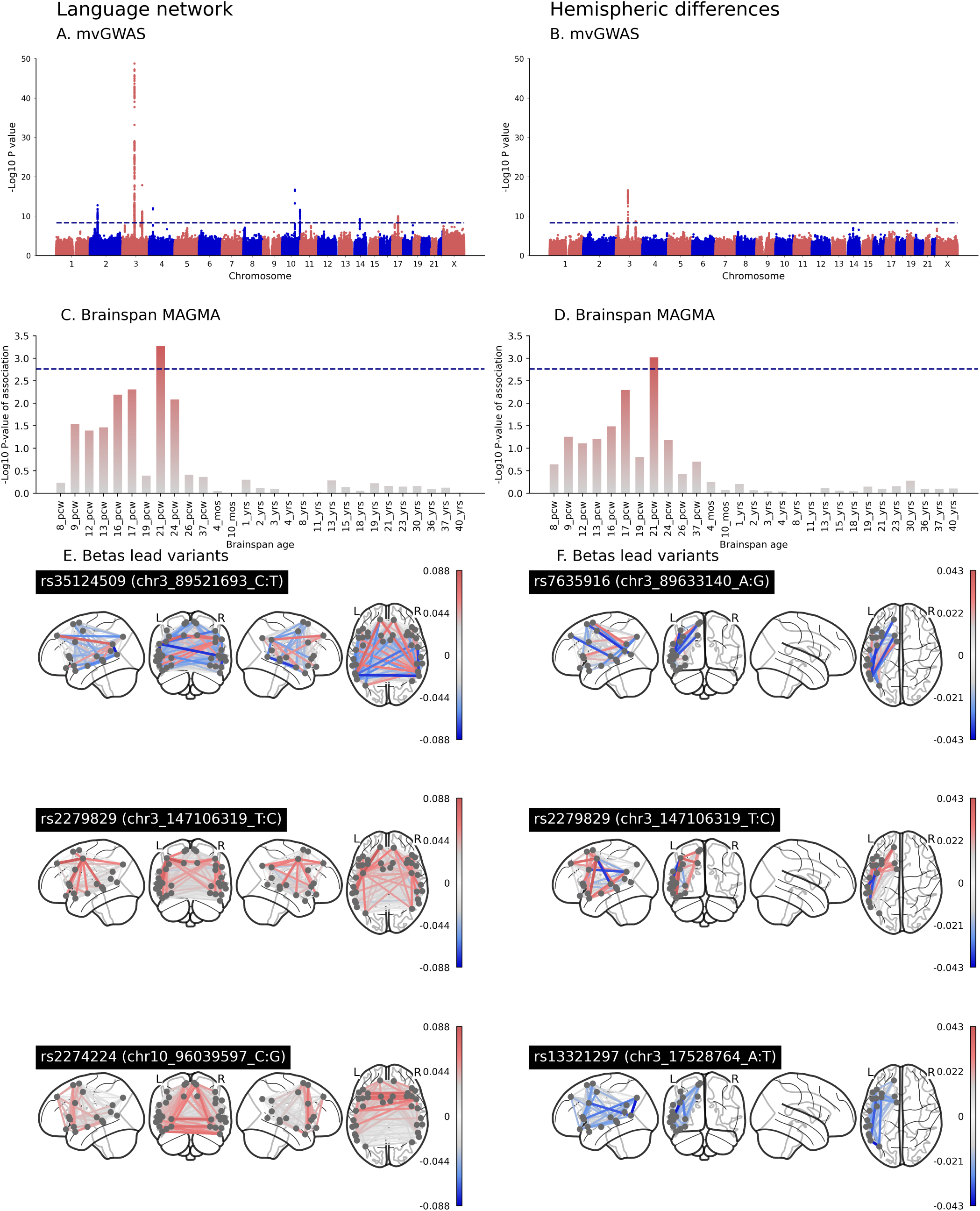
Associations with language network connectivity and asymmetry, for genetic variants with population frequencies *≥* 1 percent. **A & B**: Multivariate GWAS Manhattan plots for language network edges (A) and hemispheric differences (B). The genome is represented along the X axis of each Manhattan plot, with chromosomes in ascending numerical order and their p-to-q arms arranged from left to right. The Y axis of each Manhattan plot shows the pointwise significance of multivariate association, and each dot represents a single variant in the genome. The horizontal dashed line represents the threshold *p ≤* 5 *×* 10*^−^*^8^ for genome-wide multiple-testing correction. **C & D**: Genes associated with language network edges (C) and hemispheric differences (D) tend to be most strongly expressed in prenatal brain tissue compared to postnatal brain tissue, according to MAGMA analysis of the Brainspan gene expression database. PCW: post conception week. YRS: years. The horizontal dashed line represents the threshold for multiple testing correction across all developmental stages separately. **E & F**: Underlying univariate beta weights for the three most significant lead SNPs for language network edges, and the three most significant lead SNPs for hemispheric differences (E and F). Red indicates a positive association of a given edge or hemispheric difference with increasing numbers of the minor allele of the genetic variant, and blue indicates a negative association. Plots for all lead SNPs can be found in figure S8.

To probe the genetic effects on language network connectivity of our lead multivariate findings, we plotted the underlying univariate beta effect estimates across connectivity measures for each of the 14 lead SNPs (1E, figure S8). These showed heterogeneous effects on language network connectivity (1E). For example, lead SNP rs35124509 of the most significantly associated genomic locus on chromosome 3 was an exonic SNP in the *EPHA3* gene, where minor allele carriers (C, minor allele frequency (MAF) = 0.39) had on average generally reduced connectivity, i.e. lower time series correlations between regions, compared to non-carriers (1E, figure S5, figure S8). However, connectivity could also be higher on average for a minority of edges in these variant carriers (1E, figure S7). For the second most significantly associated genomic locus, minor allele carriers (T, MAF = 0.21) of lead SNP rs2279829 (on chromosome 3) displayed increased connectivity on average compared to non-carriers (1E, figure S6, figure S8). This SNP was located upstream from the *ZIC4* gene (figure S5). Lead SNP rs2274224 of the third most significantly associated genomic locus (on chromosome 10) is an exonic SNP in *PLCCE1:PLCE1-AS1*, (figure S7-8). That SNP showed an increase especially in interhemispheric connectivity in minor allele carriers (C, MAF = 0.44) compared to non-carriers (1E, figure S7-8). Brain spatial pattern plots for all 14 lead SNPs can be found in figure S8.

Separately, 103 hemispheric differences were also entered into a single mvGWAS, using the same procedure as for the language network edges. Three indepedent genomic loci were significantly associated with hemispheric differences (1B, table S5-6, figure S9), all of which were located on chromosome 3, and had also shown significant associations in the mvGWAS of language network edges. Lead SNP rs7625916, a different SNP in the same broader locus that encompasses *EPHA3*, showed a broadly rightward shift in hemispheric differences for carriers of the minor allele (A, MAF = 0.40), although not for all hemispheric differences (1F). This SNP was located in an intergenic region of *RP11-91A15.1* (figure S10). The lead SNP of the second locus rs2279829, located upstream of *ZIC4* was the same as for the language network edge results. Carriers of minor effect allele (C, MAF = 0.39) displayed heterogeneous changes in hemispheric differences (figure 1F, figure S11). The lead SNP for the third locus, rs13321297, located in an intronic region near *TBC1D5*, was associated with a broadly rightward shift in hemispheric differences for carriers of the minor allele (A, MAF = 0.31, figure S12). Using gene-based association mapping in FUMA we identified 9 genes associated with hemispheric differences, of which 4 were protein-coding, namely *EPHA3*, *TBC1D5*, *ZIC1* and *ZIC4*. Tissue expression of genes associated with hemispheric differences, using MAGMA as implemented in FUMA, was enriched prenatally in the Brainspan developmental data [38], reaching significance at post-conception week 21 (figure 1D). Analysis of postmortem cross-tissue expression levels from GTEx [41], and gene set analysis against 11,404 ontology and other curated sets [39, 40] , showed no significant associations after correction for multiple comparisons (figure S13 and table S7).

### Polygenic scores for language-related abilities, dyslexia and handedness

We used PRS-CS [42] to calculate genome-wide polygenic scores for language-related abilities [9], dyslexia [14] and left-handedness [19] for each of the 29,681 UK Biobank participants, using summary statistics from previous large-scale GWAS of these traits in combination with UK Biobank genotype data (Methods). Note that the previous GWAS of language-related abilities [9] was a multivariate GWAS that considered several language-related traits that had been quantitatively assessed with different neuropsychological tests: word reading, nonword reading, spelling, and phoneme awareness. After controlling for covariates (see Methods), polygenic disposition towards higher language-related abilities in the UK Biobank individuals was weakly negatively correlated with polygenic disposition towards dyslexia (*r* = *−*0.138*, p* = 3.504 *×* 10*^−^*^126^). Polygenic disposition towards left-handedness was not correlated with polygenic disposition as regards language-related abilities (*r* = *−*0.008*, p* = 0.147) or dyslexia (*r* = *−*0.005*, p* = 0.310). We then used canonical correlation analysis (CCA) in combination with permutation testing (see Methods, and figure S14 for the null distributions) to estimate overall associations of polygenic scores with language network edges and hemispheric differences. Polygenic disposition to higher language-related abilities showed a significant multivariate association with language network edges (canonical correlation r=0.160, *p* = 3*×*10*^−^*^4^) and with hemispheric differences (canonical correlation r=0.076, *p* = 9.9 *×* 10*^−^*^5^). The canonical correlation loadings showed that polygenic disposition to higher language-related abilities was most notably associated with stronger left-hemisphere connectivity, with less impact on right-hemisphere connectivity, which also meant a generally leftward shift in hemispheric differences (figure 2A).

**Figure 2:**
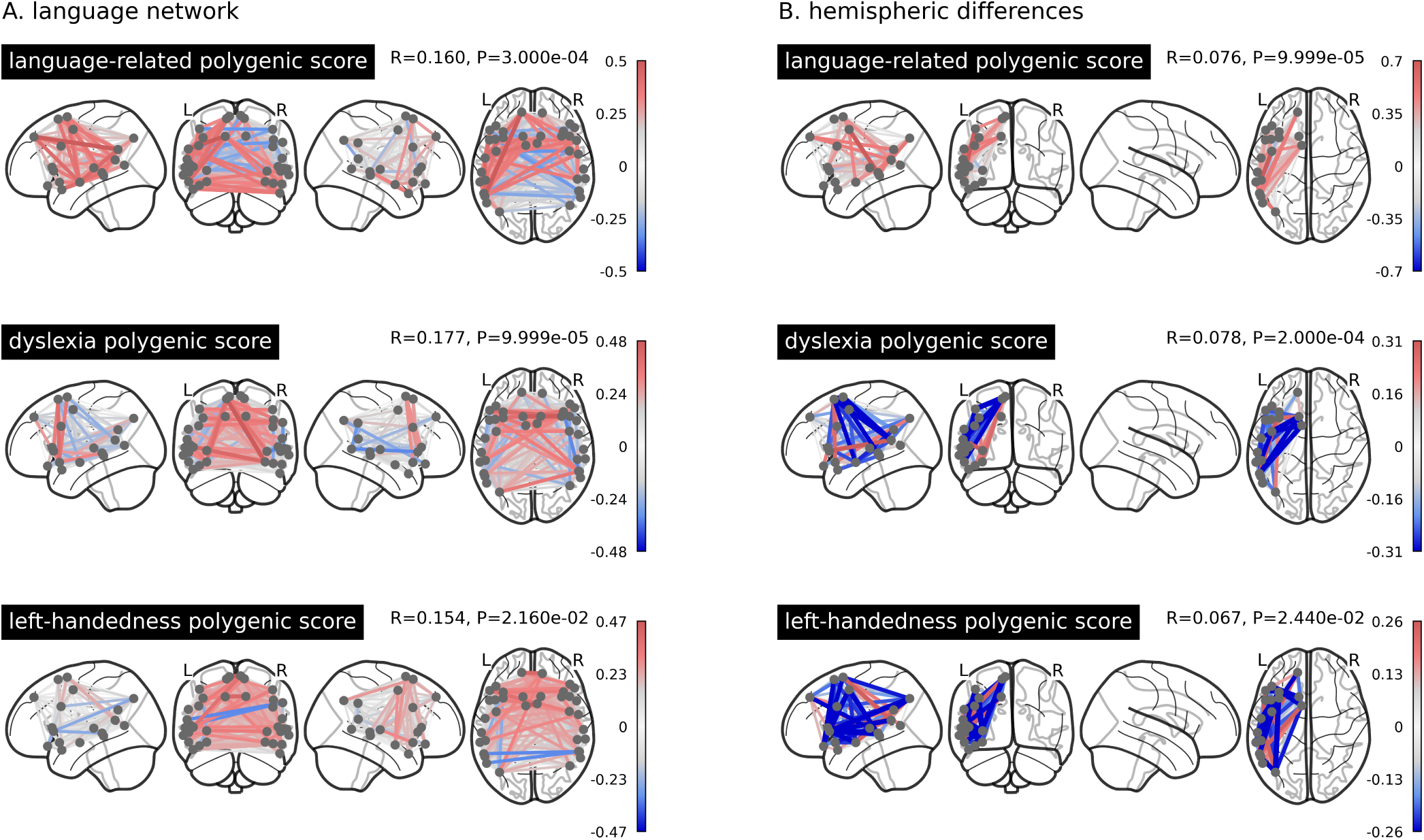
Multivariate associations with genome-wide polygenic dispositions to higher language-related abilities, dyslexia and left-handedness, for **A** the language network and **B** its hemispheric differences. Shown are the loading patterns on the first mode of six different CCA decompositions. Red indicates a positive association between polygenic score and brain phenotype, whereas blue indicates a negative association.

Polygenic disposition to dyslexia also showed significant canonical correlations with language network edges (r=0.177, *p* = 9.9 *×* 10*^−^*^5^) and hemispheric differences (r=0.078, *p* = 2 *×* 10*^−^*^4^), where especially interhemispheric connectivity was higher in those with higher polygenic disposition for this developmental reading disorder (figure 2A). In terms of hemispheric differences, higher polygenic disposition to dyslexia was associated with a broadly rightward shift in asymmetry of connectivity (figure 2B).

Polygenic disposition to left-handedness also showed significant canonical correlations: r=0.154 (*p* = 2.16 *×* 10*^−^*^2^) for language network edges and r=0.067 (*p* = 2.44 *×* 10*^−^*^2^) for hemispheric differences. Higher polygenic disposition to left-handedness was associated most notably with increased interhemispheric and right intrahemispheric connectivity, which in terms of hemispheric differences corresponds to a broadly rightward shift in asymmetry of connectivity (figure 2B).

### Rare, protein-coding variants and functional connectivity

The previous analyses were all based on genetic variants with population frequencies *>* 1 percent. We next performed a gene-based, exome-wide association scan based on protein-coding variants with frequencies *<* 1 percent, using REGENIE [43]. We used the SKAT-O gene-based test [44] for each of over 18,000 protein-coding genes with respect to 629 language network edges and 103 hemispheric differences as phenotypes, and separately using either broad (inclusive) or strict filtering for the predicted functional impacts of exonic variants (see Methods for details). Per gene we identified the lowest association *p*-value across phenotypes (Tippet’s method), and then applied an empirical exome-wide significance threshold of 2.5*×*10*^−^*^7^ to account for multiple testing across genes and phenotypes (previously established using randomized phenotypes and exome data from UK Biobank, and applied in the context of thousands of phenotypes [45]). Five genes, *NIBAN1* (*p* = 2.356 *×* 10*^−^*^7^), *MANEAL* (*p* = 1.338 *×* 10*^−^*^7^), *SLC25A48* (*p* = 4.263 *×* 10*^−^*^8^), *DUSP29* (*p* = 2.494 *×* 10*^−^*^7^) and *TRIP11* (*p* = 2.183 *×* 10*^−^*^7^), were associated with language network edges under a broad filter (figure 3A, figure S15, table S8) and 2 genes, *WDCP* (*p* = 2.064 *×* 10*^−^*^7^) and *DDX25* (*p* = 2.011 *×* 10*^−^*^8^), were associated with hemispheric differences with a strict filter (figure 3B, figure S16 and table S9).

**Figure 3:**
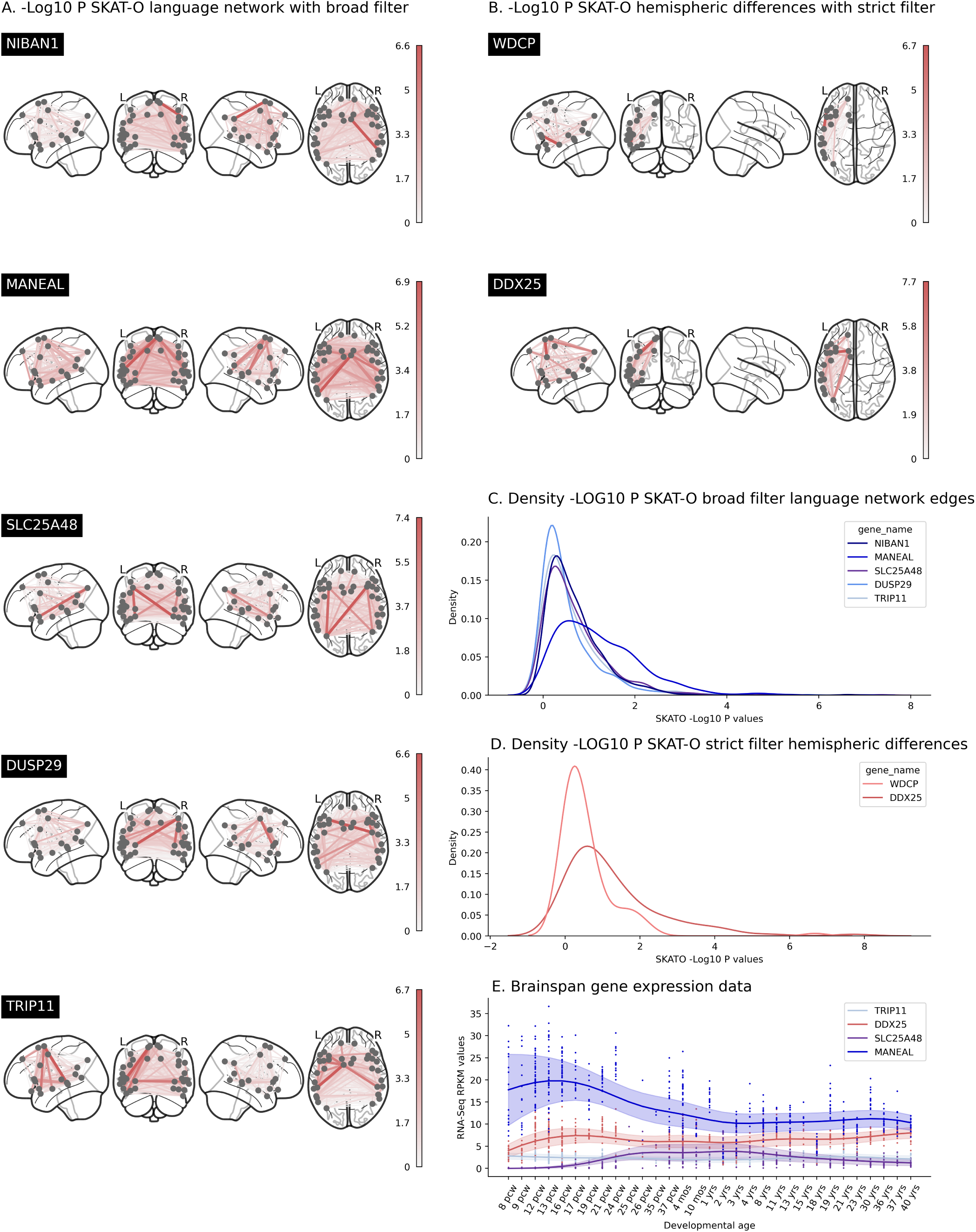
Associations of rare protein-coding variants with language network edges or hemispheric differences. **A & B.** SKAT-O -LOG10 *p*-values for genes significantly associated with the language network edges (A) and hemispheric differences (B). **C & D.** Distribution of -LOG10 *p*-values for the significantly associated genes across all brain phenotypes. **E.** RNA expression values are shown over time for all 4 genes that were available from the Brainspan dataset. Each dot represents expression levels at one timepoint in one location in the brain from one sample. Trend averages (line) and variance (shading) are shown.

For each of these 7 genes, the associations were based on multiple rare genetic variants present across multiple participants (table S10). The gene with the most distributed association pattern across functional connectivity measures of the language network was *MANEAL*, located on chromosome 1. Rare variants in this gene were most significantly associated with interhemispheric connectivity between the left middle temporal gyrus (G Temporal Mid-4-L) and the right supplementary motor area (G Supp Motor Area-3-R), with *p* = 1.34 *×* 10*^−^*^7^. SKAT-O testing is flexible for testing association when individual genetic variants might have varying directions and sizes of effects on phenotypes, but its output does not provide direct insight into these directions and effect sizes in the aggregate. We therefore followed up with a burden analysis (Methods) and found that an increased number of rare protein-coding variants in *MANEAL* was associated with generally decreased language network connectivity (figure S17).

Another gene with a distributed association pattern was *DDX25*, where rare variants were associated with multiple hemispheric difference measures. The hemispheric difference with the strongest association to this gene was between the inferior frontal sulcus (S Inf Frontal-2) and the supplementary motor area (G Supp Motor Area-2), with *p* = 2.01 *×* 10*^−^*^8^. Follow-up burden analysis showed that an increased number of *DDX25* variants that were predicted to be deleterious was associated with a broadly less leftward / more rightward shift in asymmetry (figure S18).

The five remaining genes, *NIBAN1*, *SLC25A48*, *DUSP29*, *TRIP11* and *WDCP* did not display widespread associations with respect to language network connectivity measures or hemispheric differences (figure 3C and 3D), but rather were driven by one or a few individual edges or hemispheric differences.

## Discussion

Studying the genetics of language-related brain traits, such as language network functional connectivity in the resting state, can yield clues to developmental and neurobiological mechanisms that support the brain’s functional architecture for language. In this study we report common genetic variant, polygenic and exonic rare variant associations with language network functional connectivity, and/or hemispheric differences of connectivity. We found 14 genomic loci associated with language network edges and 3 of these loci were also associated with hemispheric differences. *EPHA3* was the most significantly associated gene based on common genetic variants. A polygenic disposition for higher language-related abilities was associated with a leftward shift in functional connectivity asymmetry, while polygenic dispositions to dyslexia and left-handedness were associated with rightward shifts in functional connectivity asymmetry. Lastly, exome-wide scanning suggested 5 genes associated with language network edges and 2 genes with hemispheric differences on the basis of rare, protein-coding variants. *MANEAL* and *DDX25* showed distributed association profiles across multiple regional brain connectivity measures.

### Common variant associations

The most significant association we found was on the 3p11.1 locus, near the *EPHA3* gene, which codes for ephrin type-A receptor 3. *EPHA3* is involved in developmental processes such as neurogenesis, neural crest cell migration, axon guidance and fasciculation [46–48] and is preferentially expressed 8 to 24 weeks post-conception. This genomic locus has previously shown association with individual differences in both resting state functional connectivity [27, 28, 49] and white matter connectivity [28, 50] in the frontotemporal semantic network. Here we add to the literature that this locus is also associated with hemispheric differences of language network functional connectivity. *EPHA3* may therefore be involved in development of a left-right axis in the brain that supports hemispheric specialization for language.

A second locus associated with language network connectivity and asymmetry was located in 3p24.3, near the *TBC1D5* gene, which codes for subunit TBC1 domain family member 5. This gene may act as a GTPase-activating protein for Rab family protein(s), and is expressed in all tissues, including the brain [51]. *TBC1D5* is involved in cell processes related to macroautophagy and receptor metabolism. Recent studies have found associations of this gene with functional language network connectivity [28], white matter [52], dyslexia [14], and health-related associations with Parkinson’s Disease [53] and schizophrenia [54]. Again, here we add an association with hemispheric differences that implies a role in development of the left-right axis in the brain that supports language lateralization.

In total, of the 14 genomic loci we found, 12 were previously reported in other GWAS of brain traits [27, 28, 49, 50]. Two loci that have no previous literature associated with them in the GWAS Catalog [55] were a locus on the pseudo-autosomal part of the X and Y chromosome, with rs2360257 as lead SNP, and a locus on 3q22.2, with rs143322006 as lead SNP. The latter is intergenic to *EPHB1*, and therefore this novel finding underscores a potential role of ephrin receptors in functional connectivity of the brain’s language network. The well-known functions of ephrins in axon guidance for nerve fiber tract formation are likely to be relevant in this context.

The other 12 loci were found in two prior GWAS studies of functional connectivity [28, 49], both of which differed from each other and from the present study in terms of connectomic methodologies. This suggests that connectome methodological choices only partially influence the discovery of genetic loci, i.e. some genetic influences on brain functional connectivity can be relatively robustly detected across different methodological choices. Six out of 14 loci were also found in a study of the white matter connectome [50], which confirms that functional and structural connectivity have partially overlapping genetic architectures. The overlap of significant loci from the present study with those found in GWAS studies of dyslexia, language-related abilities and handedness was more limited, which may be expected given that the latter are behavioural traits rather than measures of brain structure or function. The 3p24.3 locus from the present study was found in a large GWAS for dyslexia [14], and the 17q21.31 locus was also associated with left-handedness [56].

### Associations with genetic predispositions

Genome-wide polygenic scores for language-related abilities, dyslexia, or left-handedness were significantly but subtly associated at the population-level with language network functional connectivity and asymmetry. These subject-level polygenic scores quantify the cumulative effects of common genetic variants from across the genome on a given trait. The leftward shift of asymmetry in people with polygenic dispositions to higher language-related abilities is consistent with functional asymmetry reflecting an optimal organization for language processing. Although language performance and functional language lateralization do not seem to be strongly correlated in healthy adults [57, 58], an absence of clear hemispheric language dominance has been reported to associate with slightly reduced cognitive functioning across multiple domains [59].

The rightward shift in asymmetry of language network connectivity with higher polygenic disposition to dyslexia is in line with some previous studies in smaller samples that suggested decreased left hemisphere language dominance in dyslexia, although this previous evidence was often inconsistent and inconclusive [60–63]. This association also converges in its direction with the association of *TBC1D5* with hemispheric differences described above. Our study therefore illustrates how large-scale brain imaging genetic analysis of genetic disposition to a human cognitive disorder can inform on the neurobiological correlates of the disorder, even when carried out using general population data.

The rightward shift in asymmetry of language network functional connectivity with higher polygenic scores for left-handedness that we observed is consistent with increased right hemisphere language dominance in left-handers [2, 20, 64]. Causality cannot be determined in a cross-sectional dataset of the kind used in our study. For example, genetic disposition may affect prenatal brain development in ways that alter functional asymmetries, and this seems likely given that many of the relevant genes are upregulated in the prenatal brain, and that functional asymmetries already exist in neonates [15]. However, some functional asymmetries may also follow, or be reinforced through, behaviours that are influenced by genetic disposition [19]. Consistent with this latter possibility, a meta-analysis of neuroimaging studies of dyslexia suggested that reduced left-hemisphere dominance is only present in adults and not in children [61]. The UK Biobank consists of middle-aged and older adults, but future studies of polygenic risk for dyslexia should test the association with brain connectiviy in younger samples, to help address the developmental/aging questions.

It is important to recognize that gene-brain associations in general population data are usually subtle [19, 65] and also that canonical correlations tend to increase with the number of variables, due to higher degrees of freedom [66]. However, as we only used the first canonical mode and only tested a single polygenic score on one side of the correlation in each analysis (versus multiple brain traits on the other side), then the freedom of the canonical correlation was relatively restricted. The permutation test that we used showed that all multivariate associations with polygenic scores were greater than expected by chance. Furthermore, the first canonical mode has previously been shown to be the most replicable [67] as it captures the most variance. Cross-validation in canonical correlation analysis is often employed for supervised model evaluations, but our use here was unsupervised and descriptive, for which there is no clear procedure for model evaluation [66]. Our interest was to describe the most accurate overall association between polygenic disposition to a given trait and brain functional connectivity measures in the available sample.

### Exome-wide scans

We report associations of 5 genes, *NIBAN1*, *MANEAL*, *SLC25A48*, *DUSP29* and *TRIP11*, with language network connectivity and 2 genes, *WDCP* and *DDX25*, with hemispheric differences on the basis of rare, protein-coding variants from exome sequence data. No previous rare variant associations have been reported with any of these 7 genes [32, 33], but *MANEAL* has been previously implicated in a GWAS of mathematical ability based on common genetic variants [68], which testifies broadly to its relevance for cognitive function. The protein encoded by *MANEAL* is found in the Golgi apparatus [69] and may regulate alpha-mannosidase activity. Previous work has shown relatively high expression of this gene in the brain compared to various other tissues [51]. *DDX25* is a DEAD box protein with the Asp-Glu-Ala-Asp motif, involved in RNA processing. Tissue expression for *DDX25* is also relatively high in the brain or testis compared to other tissues [51]. The roles of these 7 genes in brain development and function remain to be studied, for example using model systems such as cerebral organoids or knockout mice.

The exome-wide association analysis that we used here involved mass univariate testing with respect to brain connectivity measures, rather than multivariate modelling. For common genetic variants, several multivariate association frameworks have been developed, one of which we used here for our common variant GWAS (MOSTest) [35]. Such methods generally provide increased statistical power to detect effects compared to mass univariate testing, when genetic variants are associated with phenotypic covariance. However, such multivariate methods are currently lacking for application to the study of rare, protein-coding variants in Biobank-scale samples, where the effects of individual variants must be aggregated at the gene level and computational feasibility is an important consideration. The development of new multivariate methods for exome-wide analysis is required. As the findings in our exome-wide association scan only surpassed the multiple testing correction threshold by a small amount, we regard these findings as tentative until they might be replicated in the future in other datasets.

### Limitations

Resting state functional connectivity does not provide a direct measurement of language lateralization. In this study we quantified resting state functional connectivity between regions that were previously found to be involved in language on the basis of fMRI during sentence-level reading, listening and production tasks [3], and also where left-right homotopic regions were defined for the investigation of hemispheric differences. The use of full correlations as connectivity measures, as is common in the field, means that an increase in connectivity between a pair of regions can also be indirect through other regions [70]. Another caveat is that individual anatomical differences may seep into functional connectivity measures when a hard parcellation is used [70, 71]. However, as the literature has shown more broadly, structural brain properties can make meaningful contributions to functional connectivity and it might not be possible to fully disentangle the two [72–75].

Issues with respect to our chosen methods for genetic association testing have been discussed above. A general point is that we used one large discovery sample of 29,681 participants to maximize power in our GWAS, polygenic association analysis, and exome-wide scan. This did not allow for a discovery-replication design. However, using the largest available sample leads to the most accurate estimate of any possible association, including of its effect size. In light of this, the utility of discovery-replication designs has declined in relevance with the rise of biobank-scale data [76].

A limitation of the UK Biobank is that participation is on a voluntary basis, which has led to an overrepresentation of healthy participants rather than being fully representative of the general population [65, 77].

## Conclusion

In conclusion, we report 14 genomic loci associated with language network connectivity or its hemispheric differences based on common genetic variants. Polygenic dispositions to lower language-related abilities, dyslexia and left-handedness were associated with generally reduced leftward asymmetry of functional connectivity in the language network. Exome-wide association analysis based on rare, protein-altering variants (frequencies *≤* 1 %) suggested 7 additional genes. These findings shed new light on the genetic contributions to language network connectivity and its hemispheric differences based on both common and rare genetic variants, and reveal genetic links to language- and reading-related abilities and hemispheric dominance for hand preference.

## Materials and methods

### Participants

Imaging and genomic data were obtained from the UK Biobank [34] as part of research application 16066 from primary applicant Clyde Francks. The UK Biobank received ethical approval from the National Research Ethics Service Committee North West-Haydock (reference 11/NW/0382), and all of their procedures were performed in accordance with the World Medical Association guidelines. Informed consent was obtained for all participants [78]. Analyses were conducted on 29,681 participants that remained after quality control of genotype, exome and imaging data (see below).

### Imaging data

Brain imaging data were collected as described previously [79, 80]. In this analysis resting state fMRI data were used (UK Biobank data-field 20227, February 2020 release [79, 80]). Identical scanners and software platforms were used for data collection (Siemens 3T Skyra; software platform VD13). For collection of rs-fMRI data, participants were instructed to lie still and relaxed with their eyes fixed on a crosshair for a duration of 6 minutes. In that timeframe 490 datapoints were collected using a multiband 8 gradient echo EPI sequence with a flip angle of 52 degrees, resulting in a TR of 0.735 s with a resolution of 2.4×2.4×2.4mm^3^ and field-of-view of 88×88×64 voxels. Our study made use of pre-processed image data generated by an image-processing pipeline developed and run on behalf of UK Biobank (see details below).

### Genetic data

Genome-wide genotype data (UK Biobank data category 263) was obtained by the UK Biobank using two different genotyping arrays (for full details see [34]). Imputed array-based genotype data contained over 90 million SNPs and short insertion-deletions with their coordinates reported in human reference genome assembly GRCh37 (hg19). In downstream analyses we used both the unimputed and imputed array-based genotype data in different steps (below). Exome sequencing data were obtained and processed as described in more detail elsewhere [32, 45, 81] (UK Biobank data category 170, genome build GRCh38). Briefly, the IDT xGen Exome Research Panel v.1.0 was used to capture exomes. Samples were sequenced using the Illumina NovaSeq 6000 platform with S2 (first 50,000 samples) or S4 (remaining samples) flow cells and were processed by the UK Biobank team according to the OQFE Protocol (https://hub.docker.com/r/dnanexus/oqfe). Analyses using individual-level exome data (UK Biobank data field 23157) were conducted on the Research Analysis Platform (https://UKBiobankiobank.dnanexus.com)

### Sample-level quality control

Sample-level quality control at the phenotypic and genetic level was conducted on 40,595 participants who had imaging, genotype and exome data available. In phenotype sample-level quality control, participants were first excluded with imaging data labelled as unusable by UK Biobank quality control. Second, participants were removed based on outliers (here defined as 6 *×* interquartile range (IQR)) in at least one of the following metrics: discrepancy between rs-fMRI brain image and T1 structural brain image (UK Biobank field 25739), inverted temporal signal-to-noise ratio in preprocessed and artefact-cleaned preprocessed rs-fMRI (data fields 25743 and 25744), scanner X, Y and Z brain position (fields 25756, 25757 and 25758) or in functional connectivity asymmetries (see section* Imaging data preprocessing and phenotype derivation). Third, participants with missing data in the connectivity matrices were excluded. In total 3,472 participants were excluded in the phenotype QC.

Subsequently, in genetic sample-level quality control, only participants in the pre-defined white British ancestry cluster were included (data-field 22006) [34], as this was the largest single cluster in terms of ancestral homogeneity – an important consideration for some of the genetic analyses that we carried out (below). Furthermore, participants were excluded when self-reported sex (data-field 31) did not match genetically inferred sex based on genotype data (data-field 22001) or exome data, when sex chromosome aneuploidy was suspected (data-field 22019), or when exclusion thresholds were exceeded in heterozygosity (*≥* 0.1903) and/or genotype missingness rate (*≥* 0.05) (data-field 22027). Finally, one random member of each pair of related participants (up to third degree, kinship coefficient *≥* 0.0442, pre-calculated by UK Biobank) was removed from the analysis. This led to the further exclusion of 7,442 participants. In total 29,681 participants were included in all further analyses.

### Imaging data preprocessing and phenotype derivation

Preprocessing was conducted by the UK Biobank and consisted of motion correction using MCFlirt [82], intensity normalization, high-pass filtering to remove temporal drift (sigma=50.0s), unwarping using fieldmaps and gradient distortion correction. Structured scanner and movement artefacts were removed using ICA-FIX. [83–85] Preprocessed data were registered to a common reference template in order to make analyses comparable (the 6th generation nonlinear MNI152 space, http://www.bic.mni.mcgill.ca/ServicesAtlases/ICBM152NLin6).

On the local compute cluster at the MPI for Psycholinguistics, network connectivity was derived based on the AICHA atlas [30]. Key properties of the AICHA atlas are its homotopies. For each of the 192 parcels left and right hemisphere functional homotopies were defined. Of these 192 pairs, 7 regions were previously excluded from the atlas due to poor signal on the outside of the brain [30], leaving 185 parcel pairs. Time courses were extracted from the AICHA atlas using invwarp and applywarp from FSL (v. 5.0.10 [86]) and mri segstats from Freesurfer (v.6.0.0 [87]). Correlations between time courses were derived with numpy (v.1.13.1) using Python 2.7 and were transformed to z-scores using a Fisher transform in order to achieve normality. In addition, only the upper diagonal values were used. These values can be considered a measure of connection strength between two regions. Functional hemispheric differences (L-R) were derived for each connection, and outliers (6*×* IQR) were excluded. Previous work identified 18 regions as part of the core language network in multiple language processing domains (reading, listening and speaking [3]). These 18 regions and their homotopies were used in this analysis.

Two different types of imaging-derived phenotypes (IDPs) were extracted and used in genetic analyses. First, all 630 Z-transformed correlation values were included, including both intra- and interhemispheric connectivity. Second, for all intrahemispheric connectivity edges, hemispheric differences (L-R) were included, yielding 153 edge hemispheric differences. In total this yielded 783 new IDPs for further analysis.

### Genetic variant-level QC

Four different genetic datasets were prepared, as needed for four different analysis processes:

1. Array-based genotype data were filtered, maintaining variants with linkage disequilibrium (LD) *≤* 0.9, minor allele frequency (MAF) *≥* 0.01, Hardy-Weinberg Equilibrium test *p*-value *≥* 1 *×* 10*^−^*^15^ (see [43]), and genotype missingness *≤* 0.01 for REGENIE step 1 (below). 2. Imputed genotype data were filtered, maintaining bi-allelic variants with an imputation quality *≥* 0.7, Hardy-Weinberg Equilibrium test *p*-value *≥* 1 *×* 10*^−^*^7^ and genotype missingness *≥* 0.05 for association testing in MOSTest (below). 3. For genetic relationship matrices SNPs were only used if they were bi-allelic, had a genotype missingness rate *≤* 0.02, a Hardy Weinberg Equilibrium *p*-value *≥* 1*×*10*^−^*^6^, an imputation INFO score *≥* 0.9, a MAF *≥* 0.01, and a MAF difference *≤* 0.2 between the imaging subset and the whole UK Biobank were used. 4. For exome sequence data, only variants in the 39 Mbp exome sequencing target regions were retained (UK Biobank resource 3803), excluding variants in 100 bp flanking regions for which reads were not checked for coverage and quality standards in the exome processing pipeline. Monoallelic variants (marked with a ’MONOALLELIC’ filter flag) were also removed. Then, individual-level genotypes were set to no-call if the read depth was *≤* 7 (for single nucleotide variants) or *≤* 10 (for indel variant sites) and/or if the genotype quality was *≤* 20. Variant-level filtering comprised removal of variants sites with an average GQ (which is the Phred-scaled probability that the call is incorrect) across genotypes *≤* 35, variant missingness rate *≥* 0.10, minor allele count (MAC) *≤* 1, and/or low allele balance (only for variants with exclusively heterozygous genotype carriers; *≤* 0.15 for SNV sites, *≤* 0.20 for INDEL variant sites). Transition-transversion ratios were calculated prior to and after variant-level filtering as an indicator of data quality. Filtered pVCF files were converted to PLINK binary format, dropping multi-allelic variants, and then merged per chromosome. For the X chromosome, pseudo-autosomal regions (PAR1: start - base pair 2781479, PAR2: base pair 155701383 – end, genome build GRCh38) were split off from the rest of chromosome X. Any heterozygous haploid genotypes in the non-PAR chr X were set to missing.

### Heritability analysis

Genetic relationship matrices (GRMs) were computed for the study sample. In addition to previous sample-level quality control, individuals with a genotyping rate *≤* 0.98 and one random individual per pair with a kinship coefficient *≥* 0.025 derived from the GRM were excluded from this particular analysis. For all individuals that passed quality control and heritability of each of the 783 newly derived IDPs was estimated using genome-based restricted maximum likelihood (GREML) in GCTA v. 1.93.0beta [88]. Phenotypes that passed a nominal significance heritability filter of p *≤* 0.05 were included in further analysis.

### Common variant association testing

Multivariate common variant association testing (mvGWAS) was performed using the MOSTest toolbox [35] for all heritable measures separately for all 629 heritable language network edges and all 103 heritable hemispheric differences. MOSTest fully accounts for the multivariate nature by estimating the correlation structure on permuted genotype data and then computing the Mahalanobis norm as the sum of squared de-correlated z-values across univariate GWAS summary statistics and then fitting a null distribution using a gamma cumulative density function to extrapolate beyond the permuted data to significant findings. The multivariate z-statistic from MOSTest is always positive and does not provide information on directionality. We used imputed genotype array data and the following covariates: sex, age, age^2^, age *×* sex, the first 10 genetic principle components that capture genome-wide ancestral diversity, genotype array (binary variable) and various scanner-related quality measures (scanner X, Y and Z-position, inverted temporal signal to noise ratio and mean displacement as an indication of head motion) (see table S11 for UK Biobank field IDs). For directionality purposes we used the underlying univariate associations (beta estimates) and plotted these to describe the directionality. Genome-wide significant variants were annotated using the online FUMA platform (version 1.5.2) [36]. MAGMA (version 1.08) [37] gene analysis in FUMA was used to calculate gene-based *p*-values and for gene-property analyses, to investigate potential gene sets of interest [39, 40] and to map the expression of associated genes in a tissue-specific [41] and time-specific [38] fashion. Gene sets smaller than 10 were excluded from the analysis, due to risk for statistical inflation.

### Associations with genetic predispositions

In order to understand how language network edges and hemispheric differences relate to genetic predisposition for language-related abilities (quantitatively assessed in up to 33,959 participants from the GenLang consortium) [9], dyslexia (51,800 cases and 1,087,070 controls) from 23andMe, Inc. [14] and left-handedness (33,704 cases and 272,673 controls) from UK Biobank participants without imaging data [19], we used polygenic scores and canonical correlation analysis (CCA) for each polygenic score separately. Polygenic scores were calculated with PRS-CS [42], which uses a Bayesian regression framework to infer posterior effect sizes of autosomal SNPs based on genome-wide association summary statistics. PRS-CS was applied using default parameters and a recommended global shrinkage parameter phi = 0.01, combined with LD information from the 1000 Genomes Project phase 3 European-descent reference panel. PRS-CS performed in a similar way to other polygenic scoring methods, with noticeably better out-of-sample prediction than an clumping and thresholding approach [89, 90]. Before entering polygenic scores into a CCA analysis, they were residualised for these covariates: sex, age, age^2^, age *×* sex, the first 10 genetic principle components that capture genome-wide ancestral diversity, genotype array (binary variable) and various scanner-related quality measures (scanner X, Y and Z-position, inverted temporal signal to noise ratio and mean displacement as an indication of head motion) (see table S11 for UK Biobank field IDs). Polygenic scores were then normalized using quantile transform from scikit-learn v.1.0.1 and entered into a CCA analysis, also using scikit-learn. As correlation values in CCA tend to increase with the number of variables, we permuted the polygenic scores 10,000 times to build a null distribution of correlation values between IDPs and permuted polygenic scores and tested whether the correlation values of the first mode were outside the 95th percentile of the null distribution.

### Exome-wide scan

For rare variant association testing REGENIE v.3.2.1 was used [43]. In brief, REGENIE is a two-step machine learning method that fits a whole genome regression model and uses a block-based approach for computational efficiency. In REGENIE step 1, array-based genotype data were used to estimate the polygenic signal in blocks across the genome with a two-level ridge regression cross-validation approach. The estimated predictors were combined into a single predictor, which was then decomposed into 23 per-chromosome predictors using a leave one chromosome out (LOCO) approach, with a block size of 1000, 4 threads and low-memory flag. Phenotypes were transformed to a normal distribution in both REGENIE step 1 and 2. Covariates for both steps included sex, age, age^2^, age *×* sex, the first 10 genetic principle components that capture genome-wide ancestral diversity, genotype array (binary variable) and various scanner-related quality measures (scanner X, Y and Z-position, inverted temporal signal to noise ratio and mean displacement as an indication of head motion) (see table S11 for UK Biobank field IDs). Common and rare variant association tests were run conditional upon the LOCO predictor in REGENIE step 2. Functional annotation of variants was conducted using snpEff v5.1d (build 2022-04-19) [91]. Physical position in the genome was used to assign variants to genes and were annotated with Ensembl release 105. Combined Annotation Dependent Depletion (CADD) Phred scores for variants were taken from the database for nonsynonymous functional prediction (dbNSFP) (version 4.3a) [92] using snpSift 5.1d(build 2022-04-19). Variants were then classified for downstream analysis based on their functional annotations to either be included in a ’Strict’ or ’Broad’ filter or be excluded from further analysis. The ’Strict’-filter only included variants that were annotated with a ’High’ impact on a canonical gene transcript (variant types include highly disruptive mutations like frameshifts) outside of the 5% tail end of the corresponding protein (high-impact variants in the 5% tail ends usually escape nonsense-mediated decay) or a ’Moderate’ effect on a canonical gene transcript combined with CADD Phred score *≥* 20 (these include likely deleterious protein-altering missense variants). The second ’Broad’ set of variants also included ‘High’ annotated variants affecting alternative gene transcripts outside of 5% tail ends, ‘Moderate’ annotated variants that affected canonical or alternative gene transcripts with CADD Phred scores of at least 1, and ‘Modifier’ variants that affected canonical or alternative gene transcripts with CADD Phred scores of at least 1 (see table S12). A higher CADD score entails higher predicted deleteriousness of a SNP [93]. In REGENIE step 2, we performed a gene-based SKAT-O test [44] with strict and broad variant filters based on functional annotation with all heritable IDPs. A SKAT-O test is most appropriate in our study design as we had no a priori hypothesis about the direction of the genetic effect. Multivariate exome testing was conducted separately for language network edges and hemispheric differences by using Tippet’s method which involves taking the lowest *p*-value across the phenotypes of interest. This was previously used as validation method for development of MOSTest [35] and was shown to be less sensitive than multivariate genetic association testing in common variants. We adjusted for the exome-wide gene-based multiple comparison burden using an empirical *p*-value threshold for Type 1 error control from previous work (2.5 *×* 10*^−^*^7^ [33]). This was computed as 0.05 *×* the average *p*-value from 300 random phenotypes with varying heritabilities and UK Biobank exome data and approximates 0.05 expected false positives per phenotype. We then followed up significant results using (i) burden testing for assessing the effect of genetic mutation burden on brain connectivity and (ii) confirmatory variant-level association testing on the significant genes to describe which variants drove the gene-based associations.

## Data and code availability

The primary data used in this study are from the UK Biobank. These data can be provided by UK Biobank pending scientific review and a completed material transfer agreement. Requests for the data should be submitted to the UK Biobank: https://www.ukbiobank.ac.uk. Specific UK Biobank data field codes are given in Materials and Methods. Other publicly available data sources and applications are cited in Materials and Methods. We have made our mvGWAS summary statistics available online within the GWAS catalog: https://ebi.ac.uk/gwas/. This study used openly available software and codes, specifically GCTA (https://cnsgenomics.com/software/gcta/#GREML), MOSTest (https://github.com/precimed/mostest), FUMA (https://fuma.ctglab.nl/), MAGMA (https://ctg.cncr.nl/software/magma, also implemented in FUMA), PRS-CS (https://github.com/getian107/PRScs), REGENIE (https://rgcgithub.github.io/regenie/install/) and LD score regression (https://github.com/ bulik/ldsc). Custom code for this study is available from https://github.com/jsamelink/langnet_paper. All other data needed to evaluate the conclusions in the paper are present in the paper and/or the Supplementary Materials.

## Author contributions

Conceptualization - J.S.A, M.C.P., X.Z.K., M.J., S.E.F., C.F; Methodology - J.S.A, X.Z.K., Z.S., D.S., A.C.C., B.M., S.S-N., M.J.; Software - J.S.A., X.Z.K., D.S, M.J.; Formal analysis - J.S.A., M.C.P., X.Z.K., D.S., Z.S.; Data curation - J.S.A., X.Z.K., D.S.; Writing - original draft - J.S.A.; Writing - review & editing - M.C.P., X.Z.K., A.C.C., S-S.N, Z.S., D.S., B.M., M.J., S.E.F., C.F.; Visualization - J.S.A.; Project administration - C.F.; Resources - S.E.F, C.F.; Funding acquisition - C.F., S.E.F., M.J., ; Supervision - S.E.F., C.F.

## Supporting Information Appendix (SI)

All supplementary figures and tables can be found in accompanying PDF (figures) and Excel (tables/supplementary data) files. Genome-wide multivariate summary statistics from our GWAS based on common genetic variants are available online within the GWAS catalog (https://ebi.ac.uk/gwas/).

## Acknowledgements

This research was funded by the Max Planck Society (Germany), together with grants from the Netherlands Organisation for Scientific Research (NWO) (grant number 054-15-101) and French National Research Agency (ANR, grant No. 15-HBPR-0001-03) as part of the FLAG-ERA consortium project ’MULTI-LATERAL’, a Partner Project to the European Union’s Flagship Human Brain Project, and the Language in Interaction consortium (NWO Gravitation grant number 024-001-006). The study was conducted using the UK Biobank resource under application no. 16066 with C.F. as the principal applicant. Our study made use of quality-controlled brain images generated by an image-processing pipeline developed and run on behalf of the UK Biobank. The funders had no role in study design, data collection and analysis, and the decision to publish or preparation of the manuscript. The authors thank Else Eising, Giacomo Bignardi and Tristan Looden for their thoughts on the methodology. The authors thank Fabrice Crivello and Antonietta Pepe for their involvement in the inception of this project. The authors would like to thank the research participants and employees of 23andMe, Inc. for making this work possible.

## Supplementary Information Amelink et al. 2024

**Figure S1.**
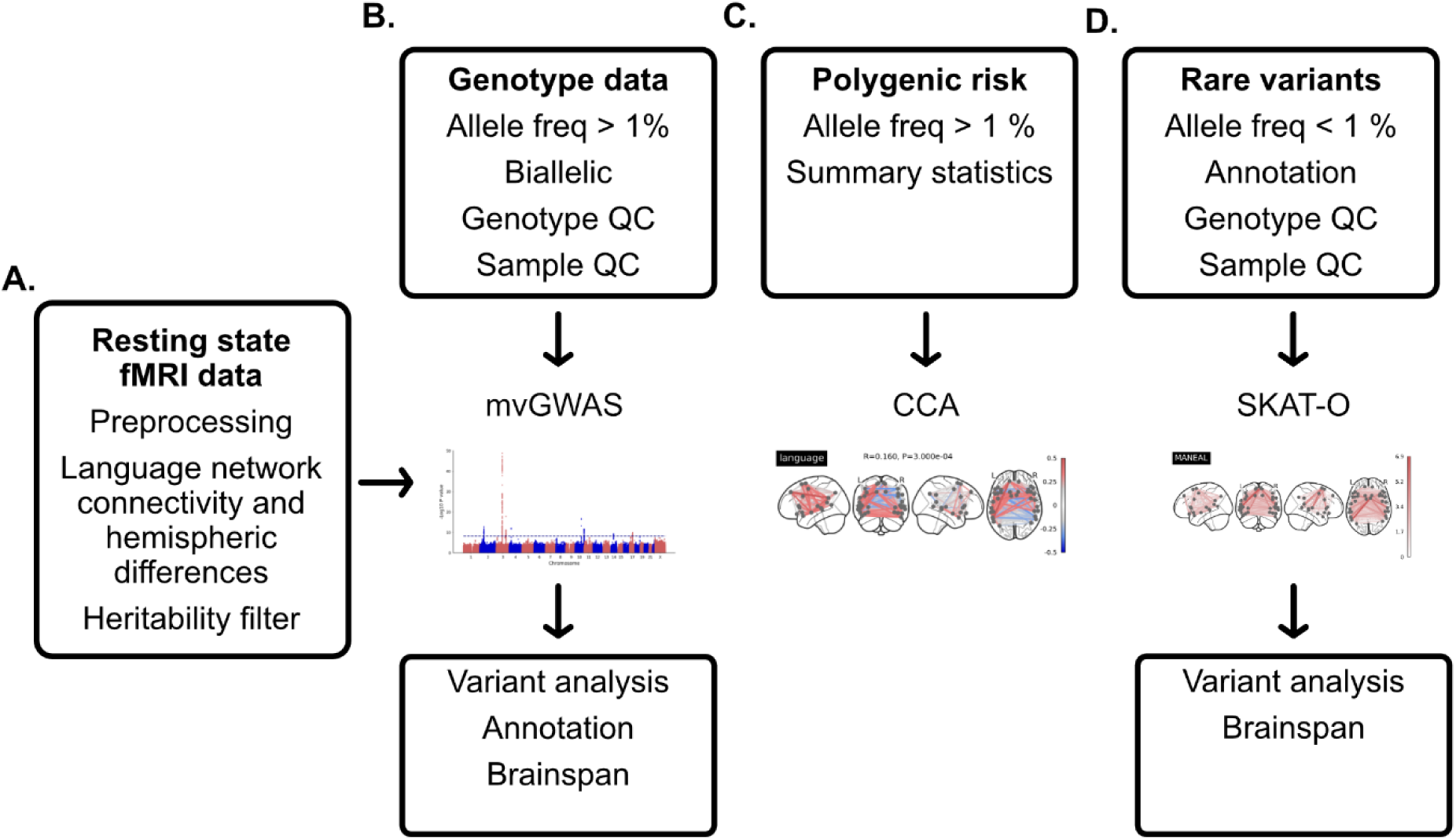
Abbreviated overview of the analysis pipelines used. **A.** We derived connectivity values and their hemispheric differences from resting state connectivity from the SENSAAS atlas that was previously developed based on several language tasks (see Introduction), filtered for heritability, and then applied three different genetic analyses to both these phenotype sets. **B.** The first analysis was a multivariate genome-wide association study (mvGWAS) based on common variant genotype data, which was annotated using FUMA, MAGMA and Brainspan data. **C.** The second analysis involved deriving of polygenic scores based on large-scale GWAS summary statistics for three phenotypes of interest: language-related performance, dyslexia, and left-handedness. We then used canonical correlation analysis (CCA) in combination with a permutation test to test the multivariate association patterns between these scores and our language network connectivity and hemispheric differences. **D.** The third analysis was an exome-wide scan using a gene-based SKAT-O test (an optimized sequence kernel association test), which was followed-up with variant association testing and annotation using Brainspan data. Allele freq represents allele frequency. Quality control is abbreviated as QC.

**Figure S2.**
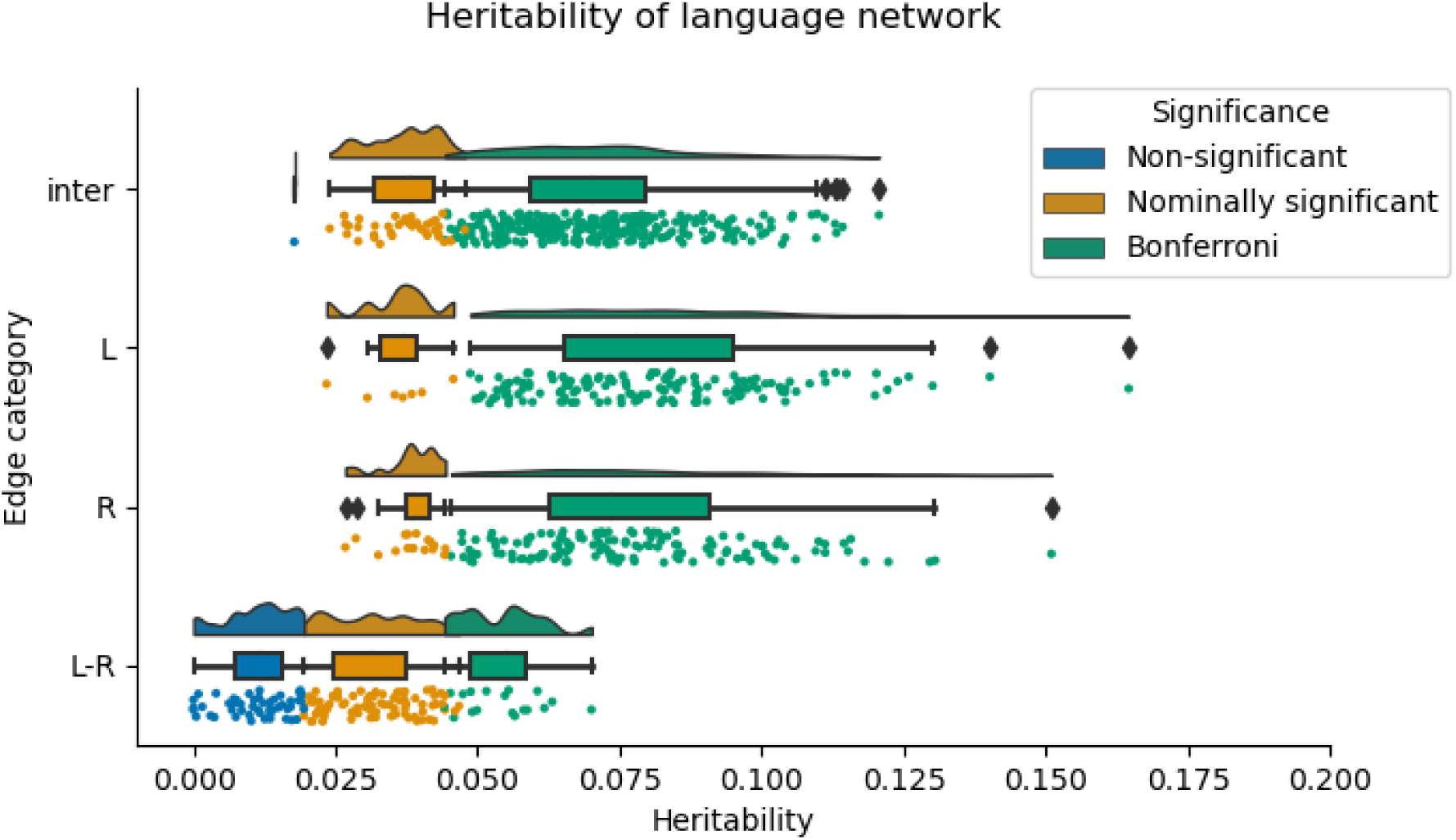
Heritability estimates from GCTA for all derived phenotypes. All non-significant phenotypes (blue) were omitted from all further analyses.

**Figure S3.**
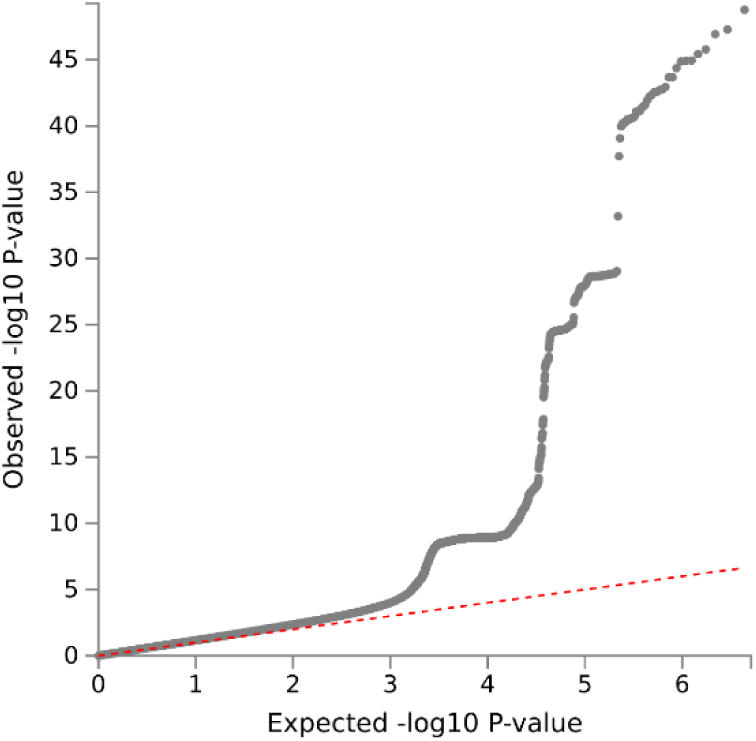
QQ plot for mvGWAS results for language network

**Figure S4.**
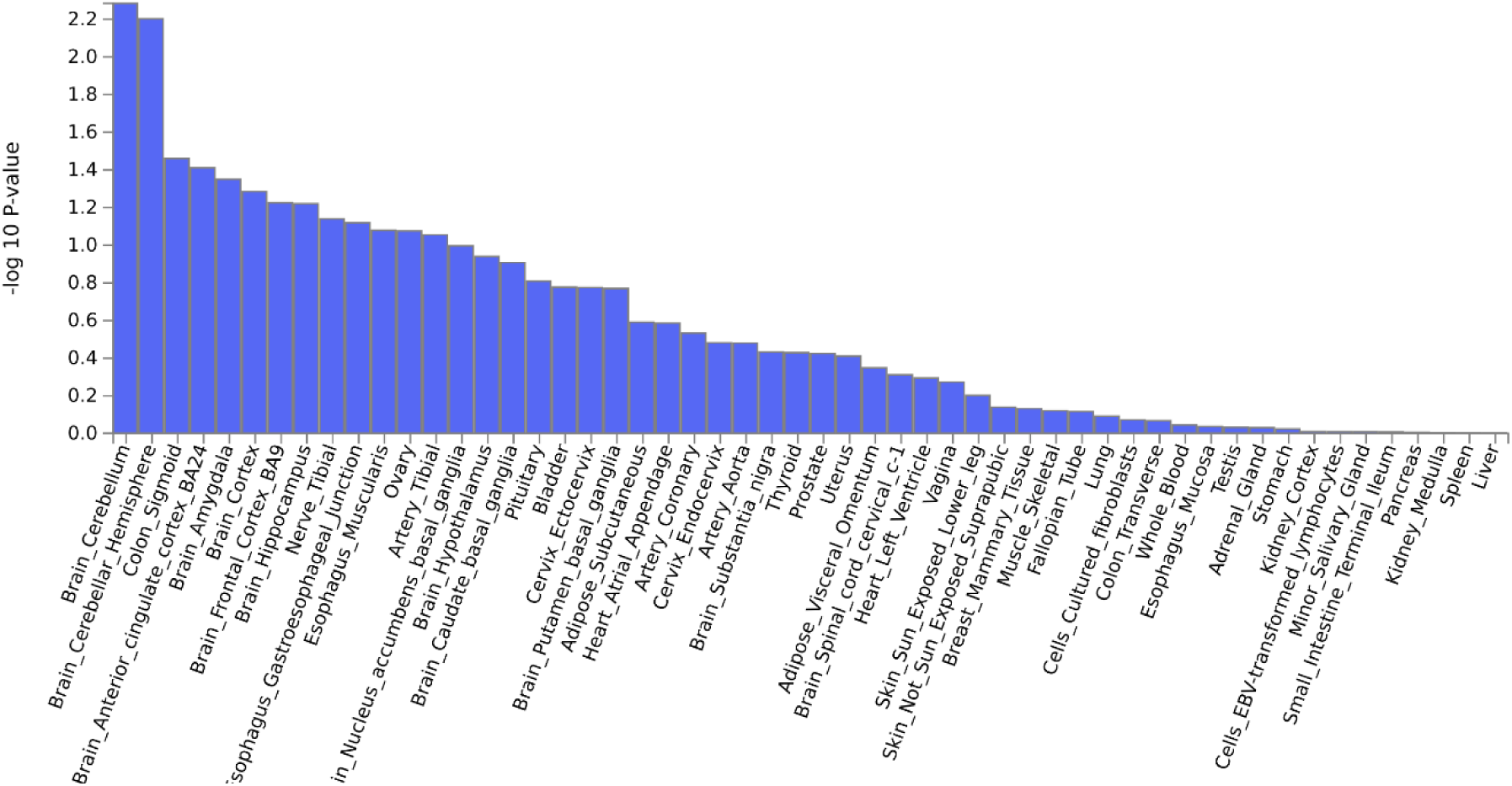
GTEx v8 53 tissue types for language network

**Figure S5.**
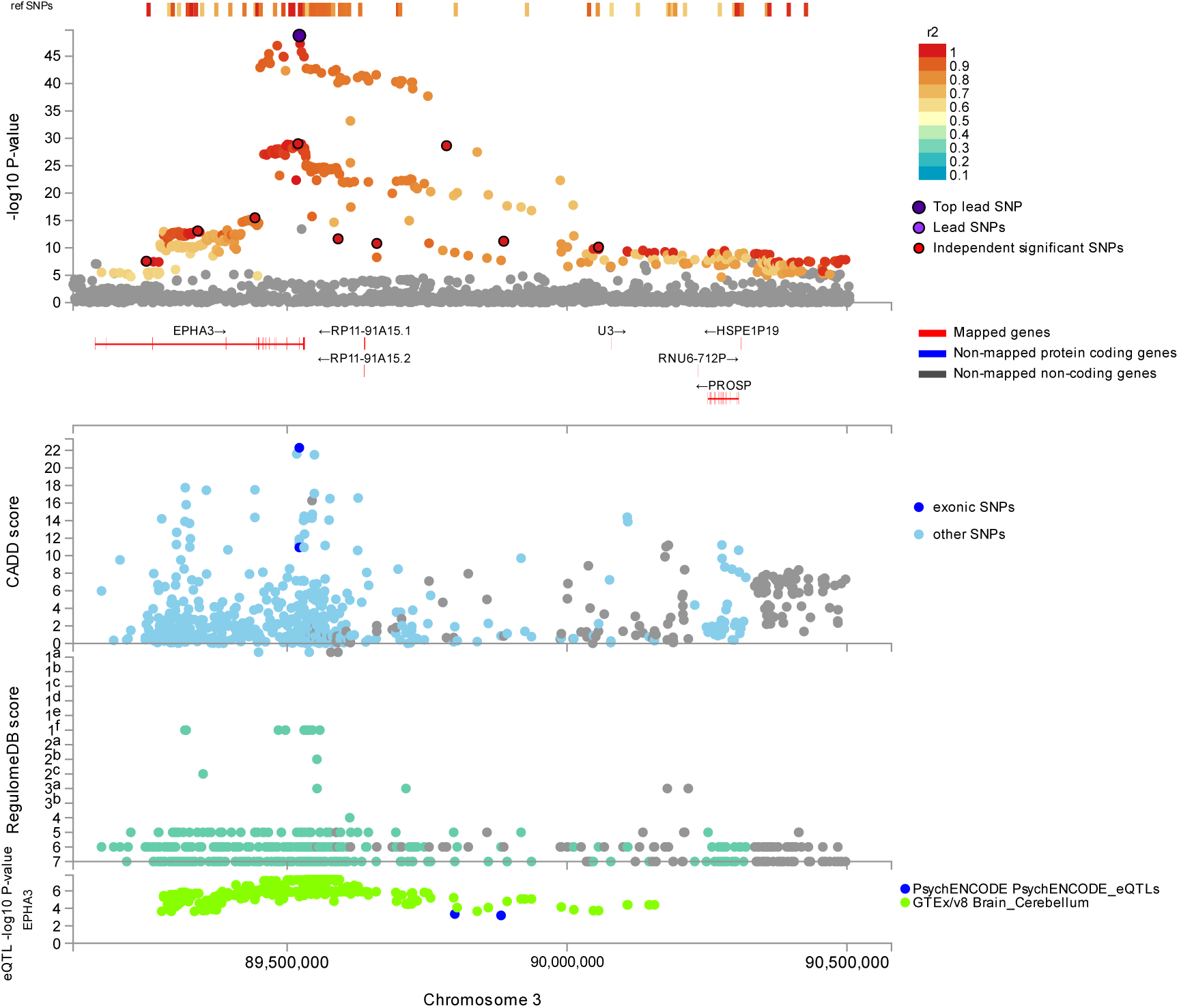
Locuszoom plot for language network results of rs35124509 on chromosome 3. Top: a fine-mapping plot is shown with lead SNPs and linkage disequilibrium (r2). Middle: Combined Annotation Dependent Depletion (CADD) scores are shown, which predict a functional protein effect. Bottom: RegulomeDB scores are shown, which predict interaction effects and gene expression effects using expression quantitative trait loci (eQTL), relating to psychiatric disorders and brain expression.

**Figure S6.**
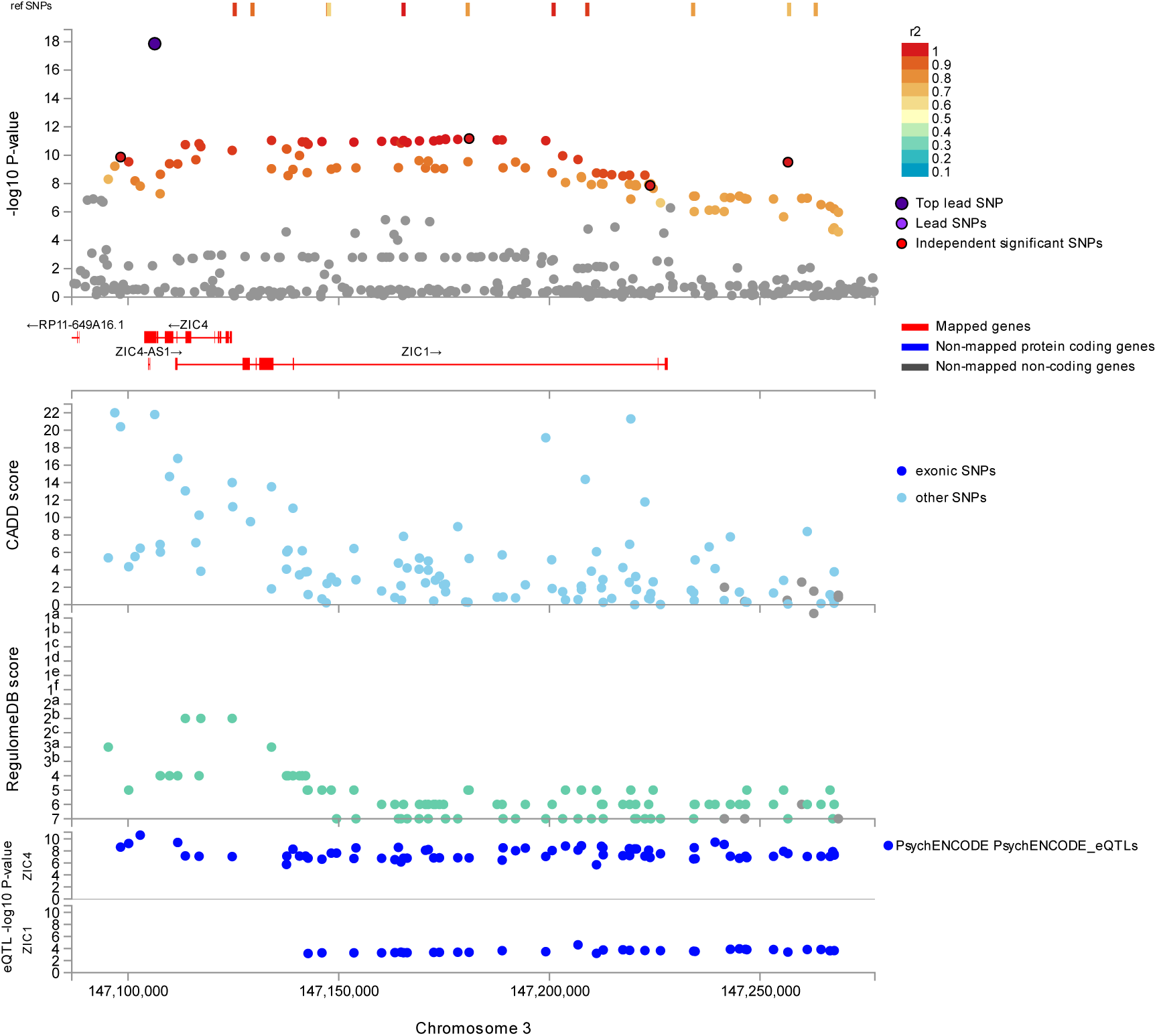
Locuszoom plot for language network results of rs2279829 on chromosome 3. Top: a fine-mapping plot is shown with lead SNPs and linkage disequilibrium (r2). Middle: Combined Annotation Dependent Depletion (CADD) scores are shown, which predict a functional protein effect. Bottom: RegulomeDB scores are shown, which predict interaction effects and gene expression effects using expression quantitative trait loci (eQTL), relating to psychiatric disorders and brain expression.

**Figure S7.**
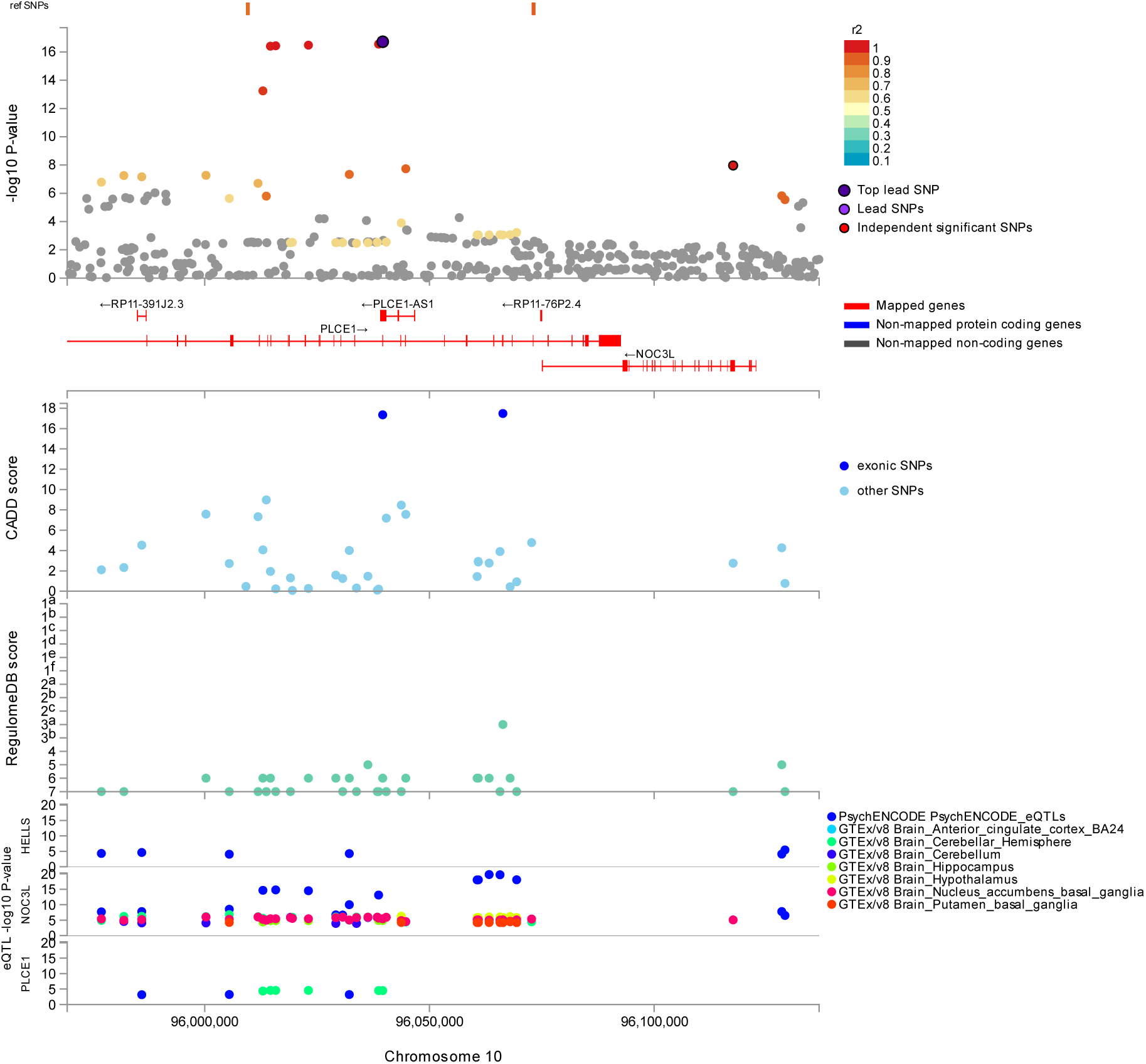
-. Locuszoom plot for language network results of rs2274224 on chromosome 10. Top: a fine-mapping plot is shown with lead SNPs and linkage disequilibrium (r2). Middle: Combined Annotation Dependent Depletion (CADD) scores are shown, which predict a functional protein effect. Bottom: RegulomeDB scores are shown, which predict interaction effects and gene expression effects using expression quantitative trait loci (eQTL), relating to psychiatric disorders and brain expression.

**Figure S8.**
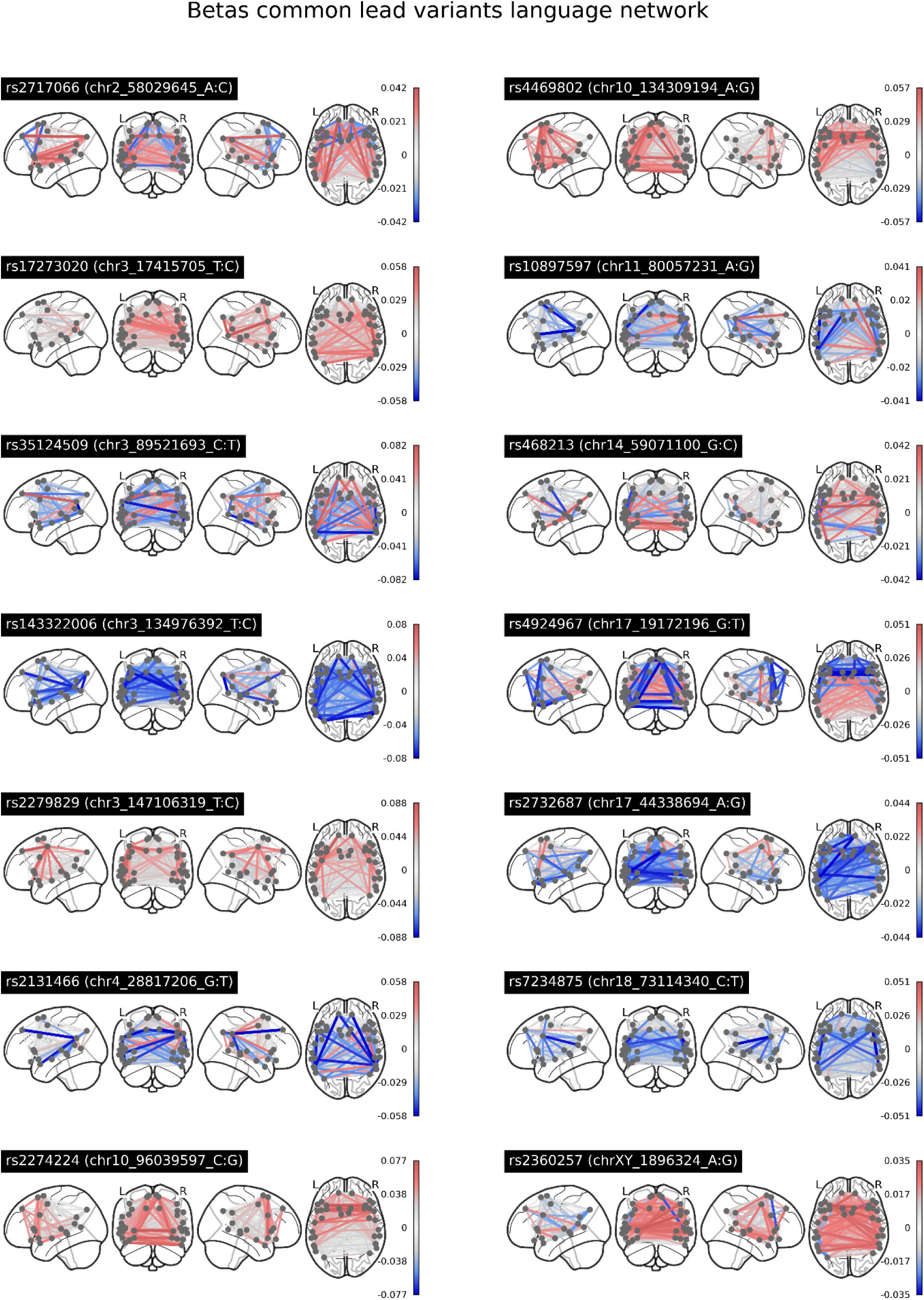
-. Underlying univariate beta weights for all 14 significant lead SNPs for language network edges. Red indicates a positive association of a given edge or hemispheric difference with increasing number of the minor allele of the genetic variant, and blue indicates a negative association.

**Figure S9.**
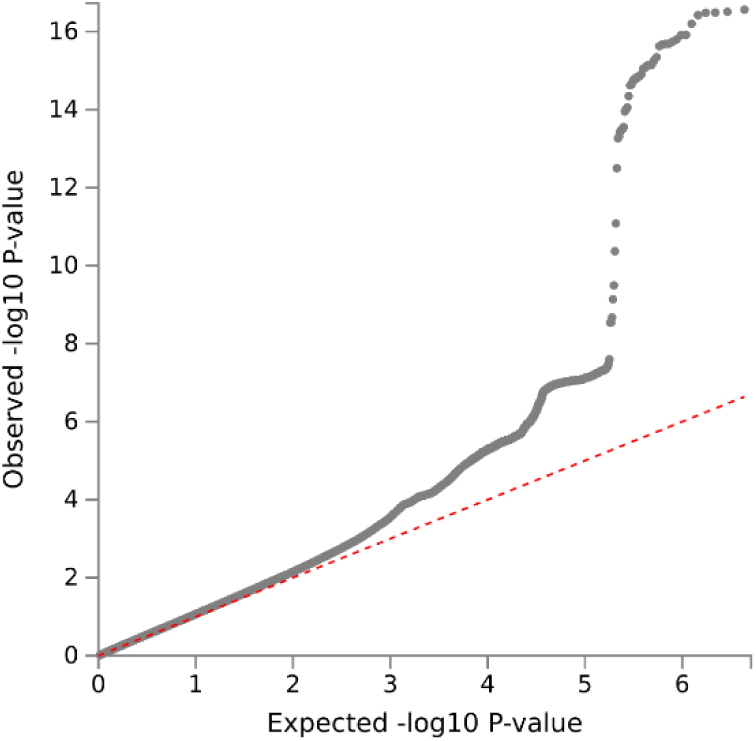
QQ plot for mvGWAS results hemispheric differences

**Figure S10.**
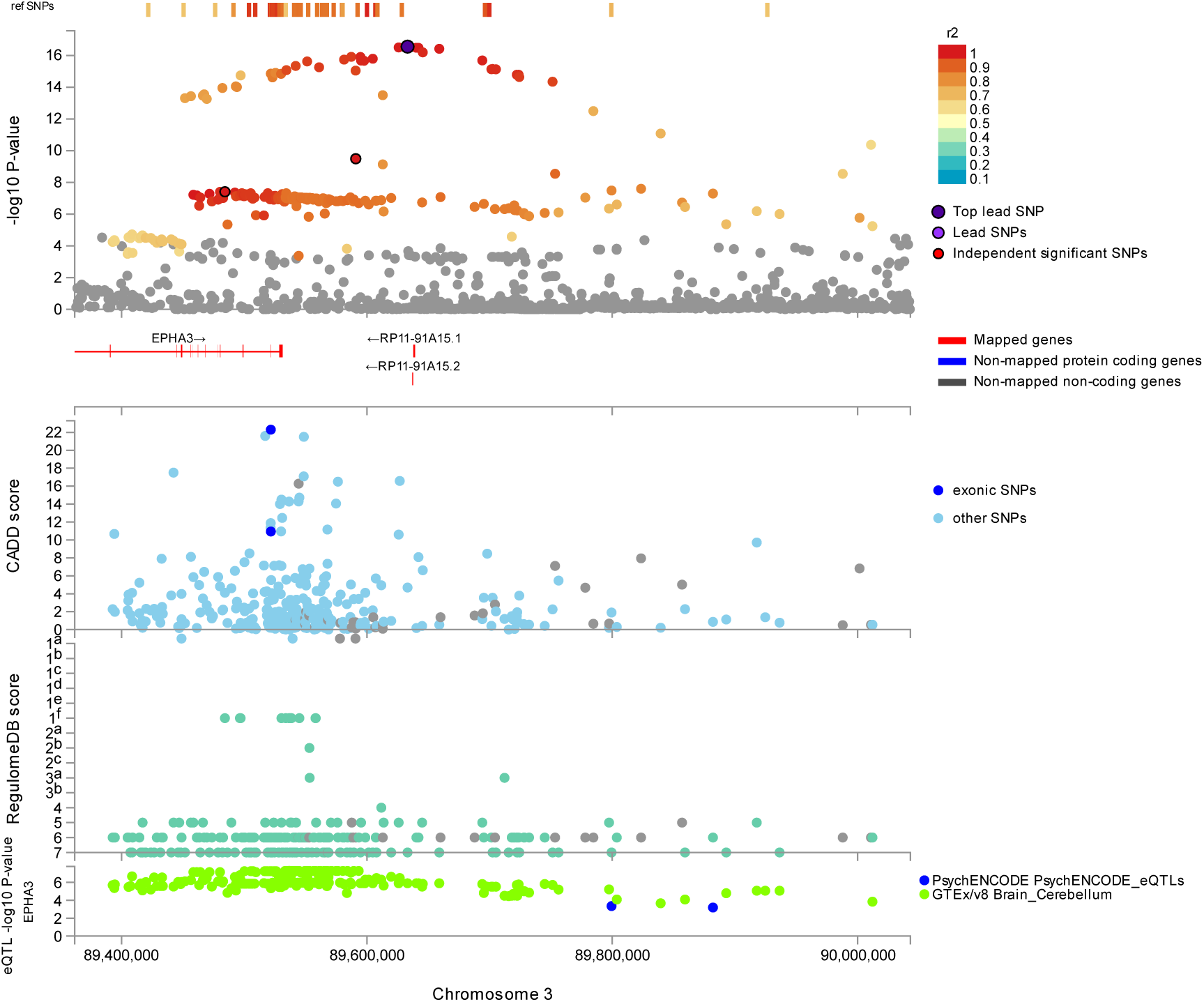
Locuszoom plot for hemispheric differences results of rs7625916 on chromosome 3. Top: a fine-mapping plot is shown with lead SNPs and linkage disequilibrium (r2). Middle: Combined Annotation Dependent Depletion (CADD) scores are shown, which predict a functional protein effect. Bottom: RegulomeDB scores are shown, which predict interaction effects and gene expression effects using expression quantitative trait loci (eQTL), relating to psychiatric disorders and brain expression.

**Figure S11.**
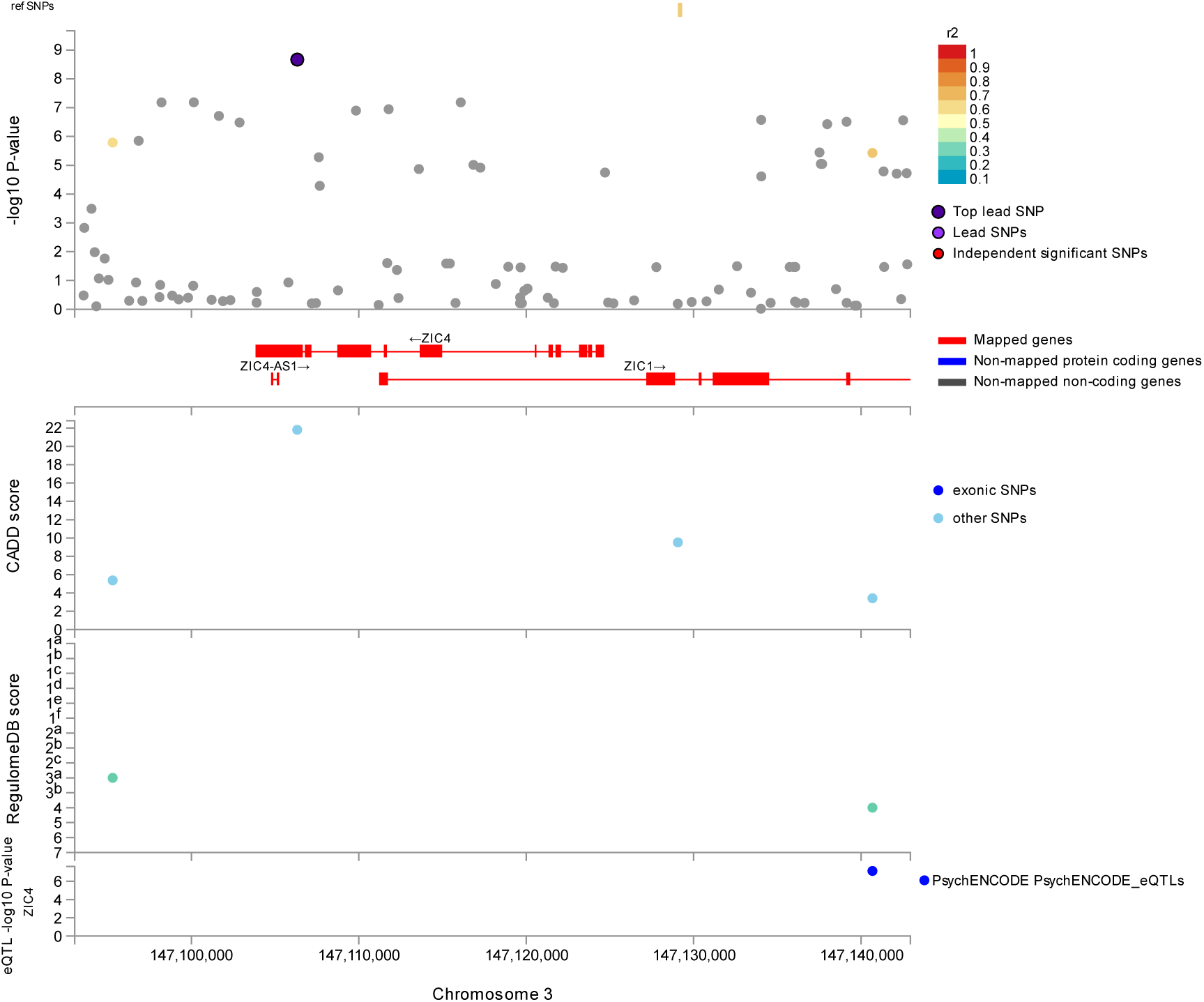
Locuszoom plot for hemispheric differences results of rs2279829 on chromosome 3. Top a fine-mapping plot is shown with lead SNPs and linkage disequilibrium (r2). Middle: Combined Annotation Dependent Depletion (CADD) scores are shown, which predict a functional protein effect. Bottom: RegulomeDB scores are shown, which predict interaction effects and gene expression effects using expression quantitative trait loci (eQTL), relating to psychiatric disorders and brain expression.

**Figure S12.**
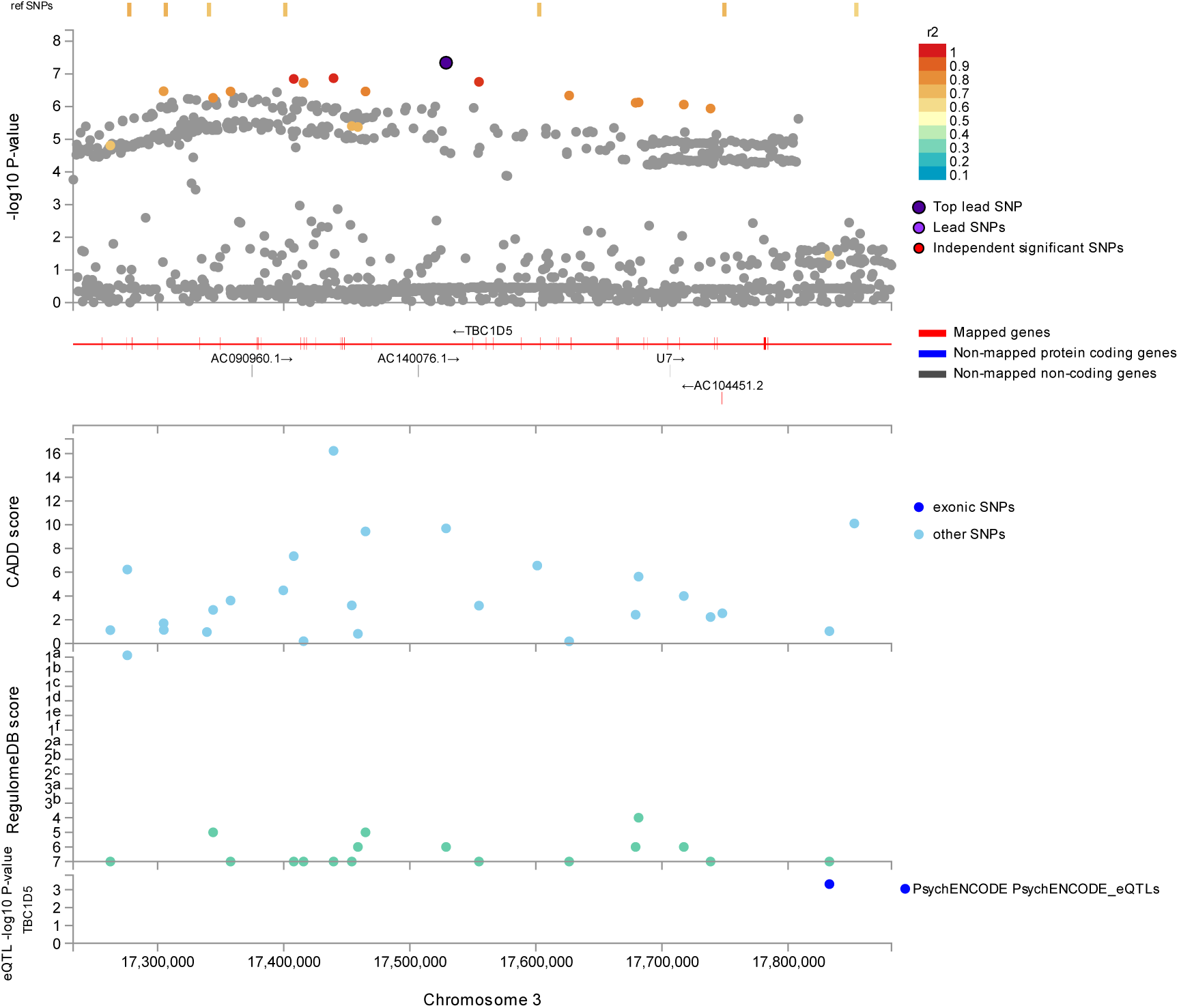
Locuszoom plot for hemispheric differences results of rs1332197 on chromosome 3. Top: a fine-mapping plot is shown with lead SNPs and linkage disequilibrium (r2). Middle: Combined Annotation Dependent Depletion (CADD) scores are shown, which predict a functional protein effect. Bottom: RegulomeDB scores are shown, which predict interaction effects and gene expression effects using expression quantitative trait loci (eQTL), relating to psychiatric disorders and brain expression.

**Figure S13.**
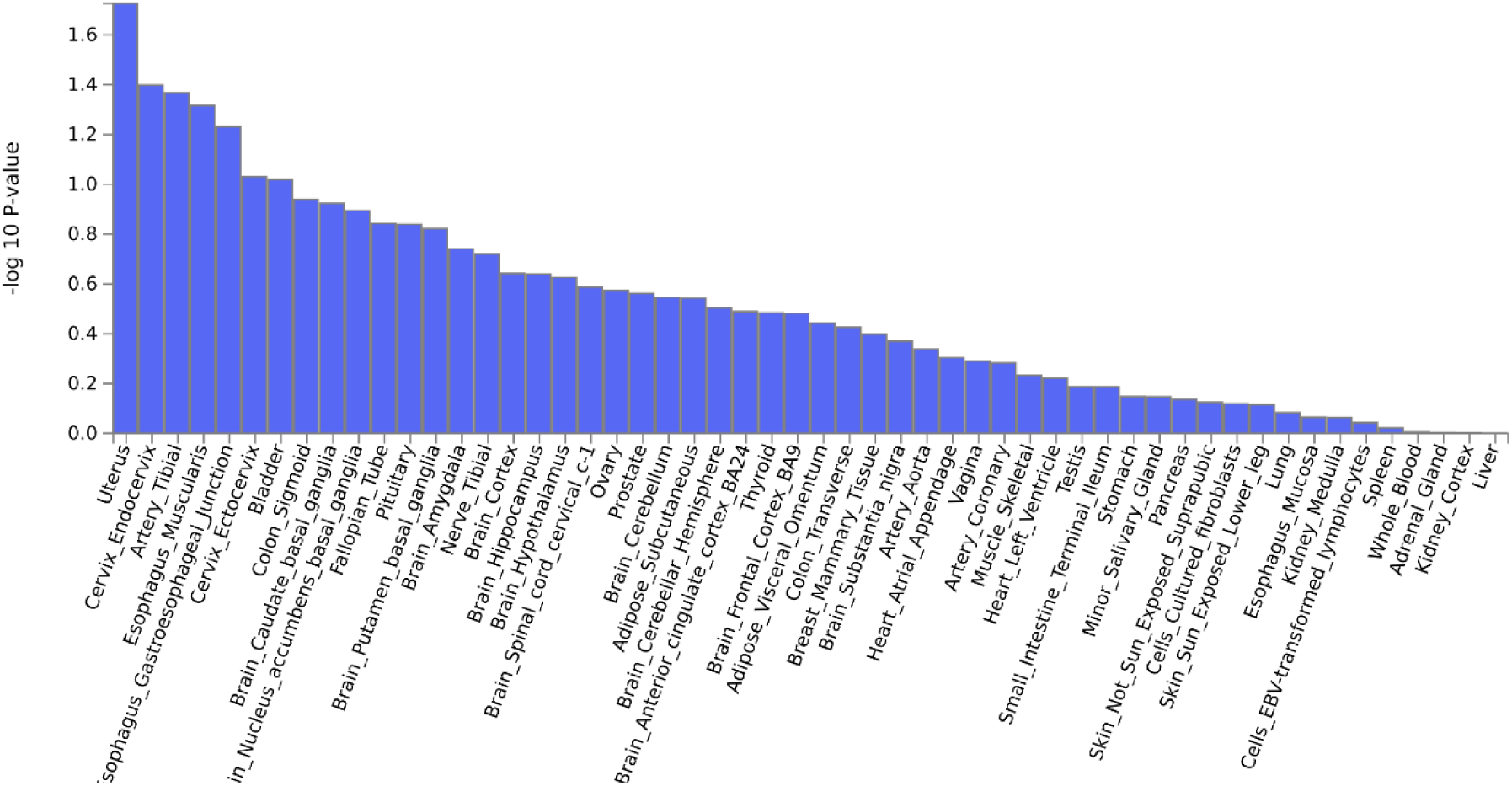
-. GTEx v8 53 tissue types for hemispheric differences

**Figure S14.**
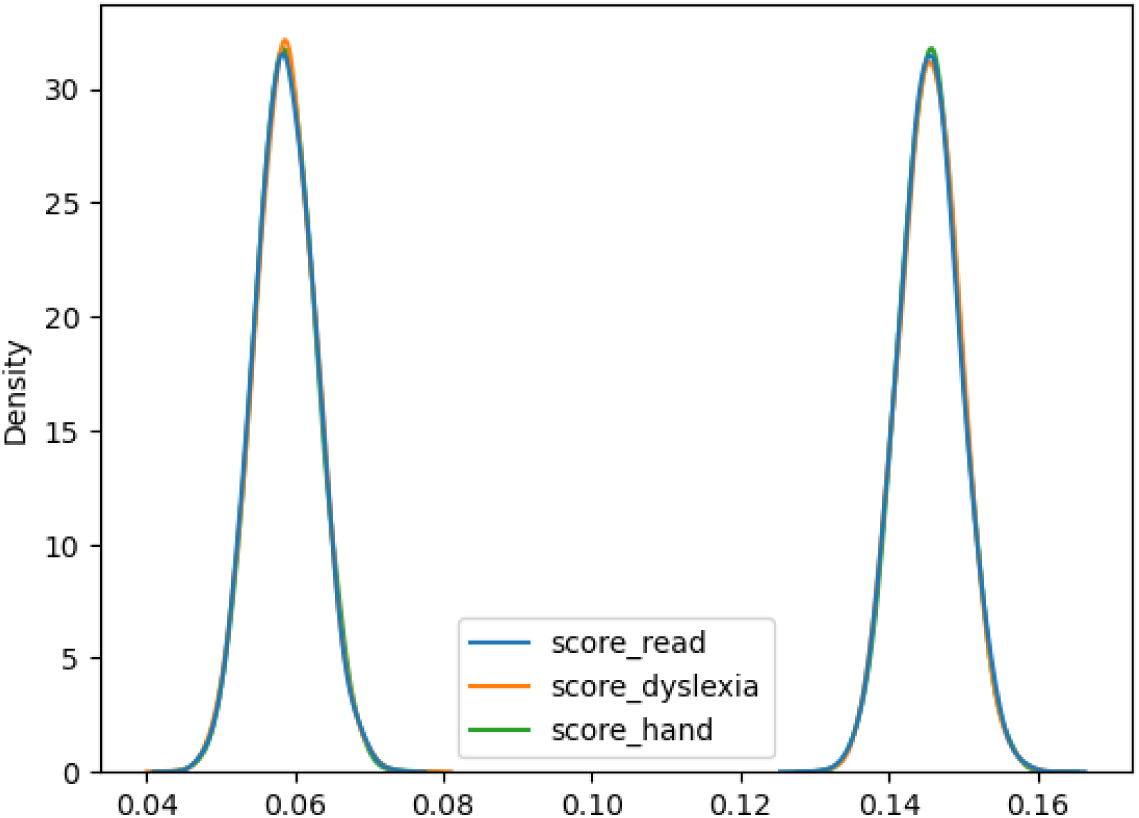
Null distributions for CCA results with permuted language-related abilities, dyslexia and left-handedness polygenic scores. Left distribution is hemispheric differences, right is language network.

**Figure S15.**
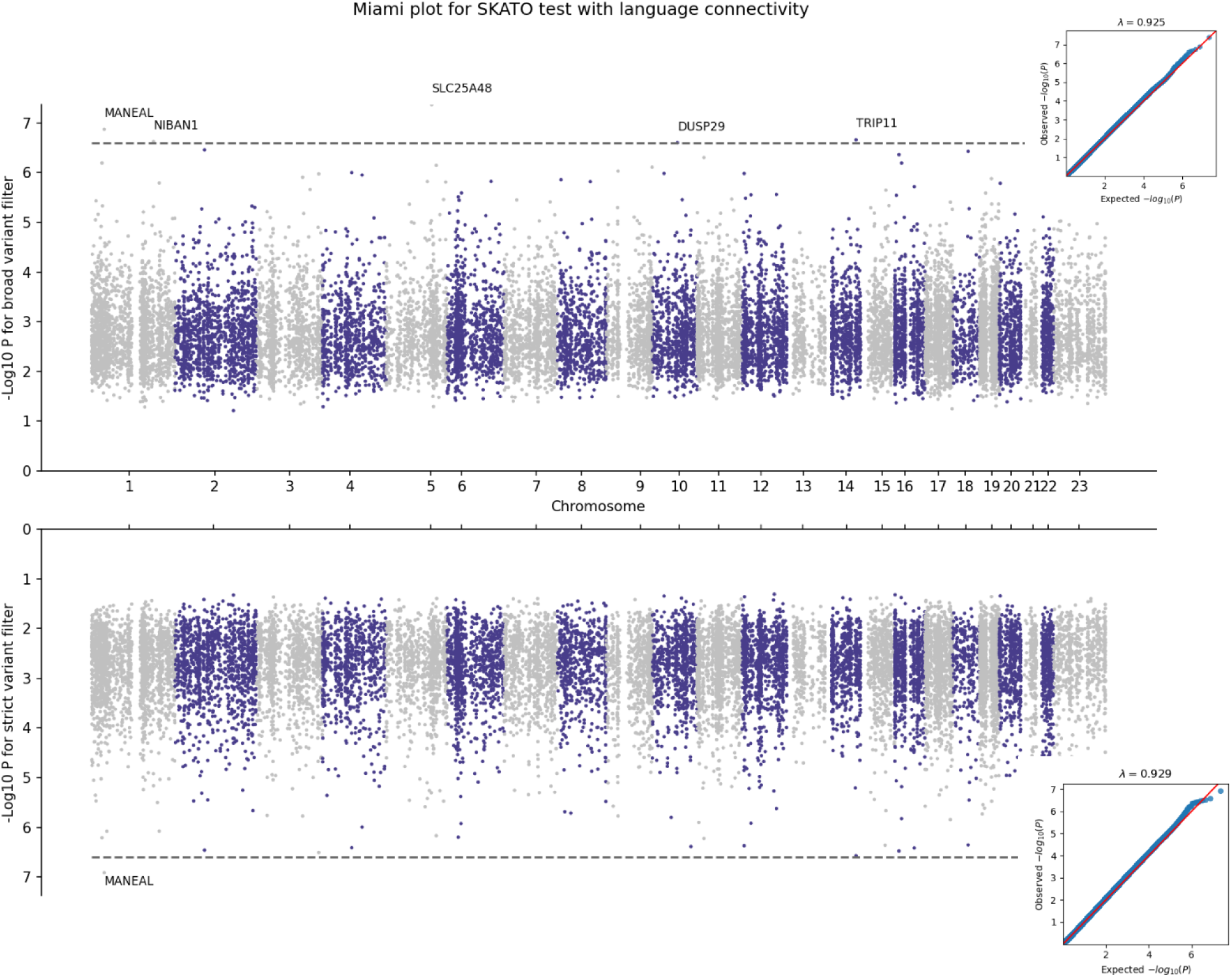
Miami plot for exome-wide gene-based lowest *p*-value associations with language network. Top results are with a broad variant filter, bottom results are with a strict variant filter. QQ plot inserts show genomic inflation for all *p*-values.

**Figure S16.**
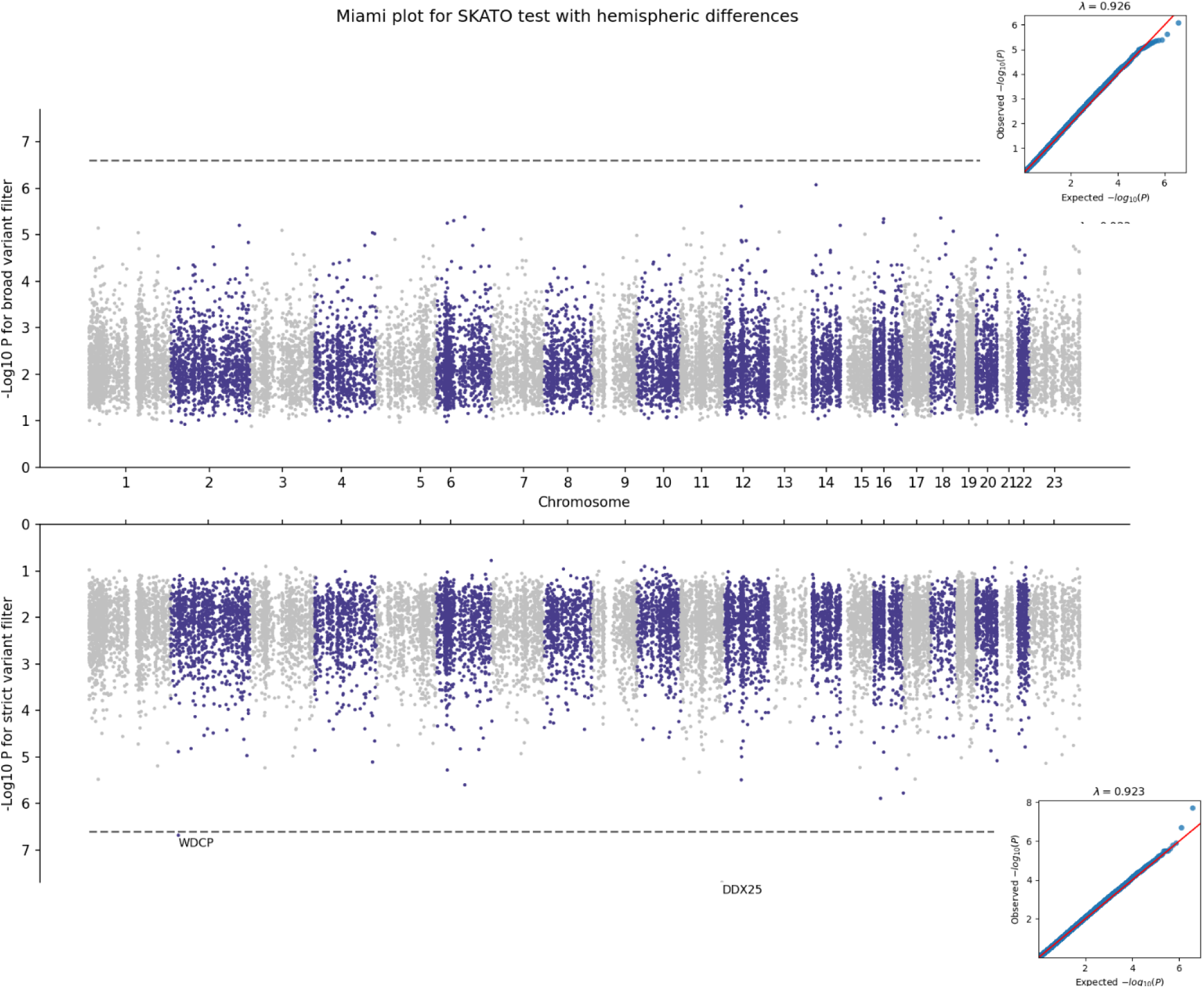
-. Miami plot for exome-wide gene-based lowest *p*-value associations with hemispheric differences. Top results are with a ‘broad’ variant filter, bottom results are with a ‘strict’ variant filter (see Methods). QQ plot inserts show genomic inflation for all *p*-values.

**Figure S17.**
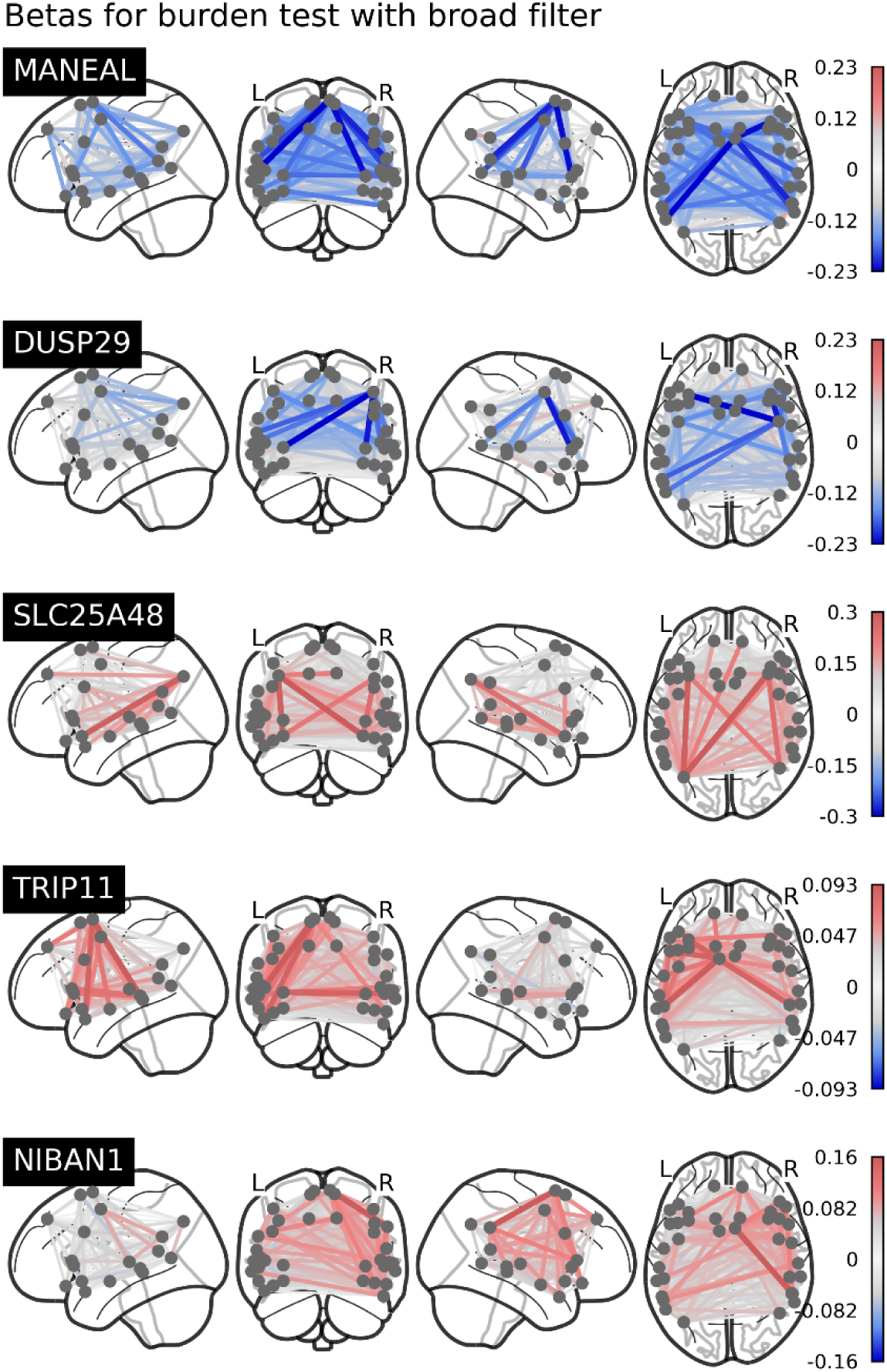
Language network betas for increased genetic burden with a ‘broad’ variant filter (see Methods). Red means an increase in connectivity, blue means a decrease in connectivity.

**Figure S18.**
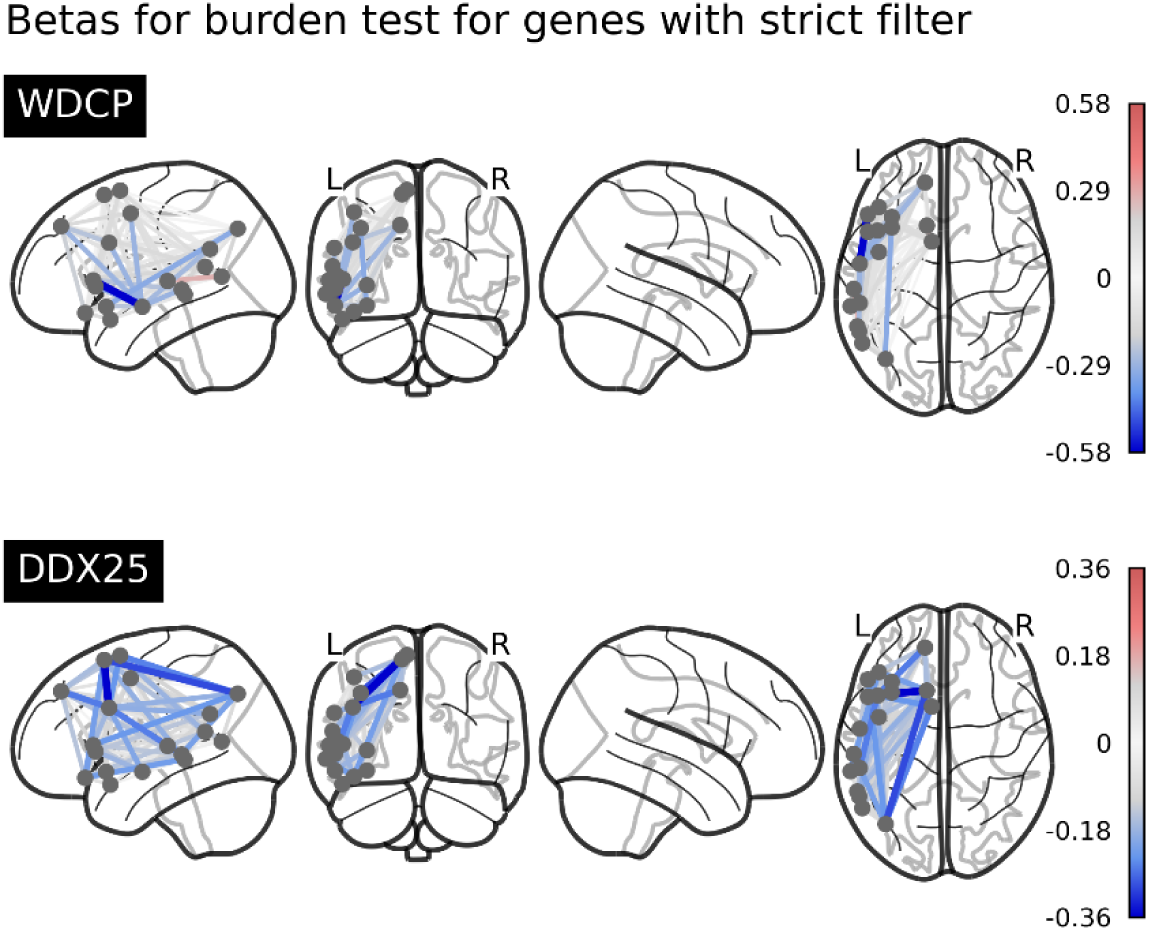
Hemispheric differences betas for increased genetic burden with a ‘strict’ variant filter (see Methods). Red means an increase in connectivity, blue means a decrease in connectivity.

## References

1. Olulade, O. A. et al. The neural basis of language development: Changes in lateralization over age. Proceedings of the National Academy of Sciences 117. Publisher: Proceedings of the National Academy of Sciences, 23477–23483. https://www.pnas.org/doi/full/10.1073/pnas.1905590117 (2023) (Sept. 2020).

2. Mazoyer, B. et al. Gaussian Mixture Modeling of Hemispheric Lateralization for Language in a Large Sample of Healthy Individuals Balanced for Handedness. en. PLOS ONE 9. Publisher: Public Library of Science, e101165. issn: 1932-6203. https://journals.plos.org/plosone/article?id=10.1371/journal.pone.0101165 (2023) (June 2014).

3. Labache, L. et al. A SENtence Supramodal Areas AtlaS (SENSAAS) based on multiple task-induced activation mapping and graph analysis of intrinsic connectivity in 144 healthy right-handers. en. Brain Structure and Function 224, 859–882. issn: 1863-2653, 1863-2661. http://link.springer.com/10.1007/s00429-018-1810-2 (2022) (Mar. 2019).

4. Malik-Moraleda, S. et al. An investigation across 45 languages and 12 language families reveals a universal language network. en. Nature Neuroscience 25. Number: 8 Publisher: Nature Publishing Group, 1014–1019. issn: 1546-1726. https://www.nature.com/articles/s41593-022-01114-5 (2023) (Aug. 2022).

5. Dale, P., et al. Genetic influence on language delay in two-year-old children. en. Nature Neuroscience 1. Number: 4 Publisher: Nature Publishing Group, 324–328. issn: 1546-1726. https://www.nature.com/articles/nn0898_324 (2023) (Aug. 1998).

6. Le Guen, Y., Amalric, M., Pinel, P., Pallier, C. & Frouin, V. Shared genetic aetiology between cognitive performance and brain activations in language and math tasks. en. Scientific Reports 8. Number: 1 Publisher: Nature Publishing Group, 17624. issn: 2045-2322. https://www.nature.com/articles/s41598-018-35665-0 (2023) (Dec. 2018).

7. Newbury, D. F., Bishop, D. V. M. & Monaco, A. P. Genetic influences on language impairment and phonological short-term memory. English. Trends in Cognitive Sciences 9. Publisher: Elsevier, 528–534. issn: 1364-6613, 1879-307X. https://www.cell.com/trends/cognitive-sciences/abstract/S1364-6613(05)00262-7 (2023) (Nov. 2005).

8. Andreola, C. et al. The heritability of reading and reading-related neurocognitive components: A multi-level meta-analysis. Neuroscience & Biobehavioral Reviews 121, 175–200. issn: 0149-7634. https://www.sciencedirect.com/science/article/pii/S0149763420306503 (2023) (Feb. 2021).

9. Eising, E. et al. Genome-wide analyses of individual differences in quantitatively assessed reading- and language-related skills in up to 34,000 people. Proceedings of the National Academy of Sciences 119. Publisher: Proceedings of the National Academy of Sciences, e2202764119. https://www.pnas.org/doi/abs/10.1073/pnas.2202764119 (2023) (Aug. 2022).

10. Verhoef, E., Shapland, C. Y., Fisher, S. E., Dale, P. S. & St Pourcain, B. The developmental origins of genetic factors influencing language and literacy: Associations with early-childhood vocabulary. en. Journal of Child Psychology and Psychiatry 62. eprint: https://onlinelibrary.wiley.com/doi/pdf/10.1111/jcpp.13327, 728–738. issn: 1469-7610. https://onlinelibrary.wiley.com/doi/abs/10.1111/jcpp.13327 (2023) (2021).

11. Eising, E. et al. A set of regulatory genes co-expressed in embryonic human brain is implicated in disrupted speech development. en. Molecular Psychiatry 24. Number: 7 Publisher: Nature Publishing Group, 1065–1078. issn: 1476-5578. https://www.nature.com/articles/s41380-018-0020-x (2023) (July 2019).

12. Deriziotis, P. & Fisher, S. E. Speech and Language: Translating the Genome. English. Trends in Genetics 33. Publisher: Elsevier, 642–656. issn: 0168-9525. https://www.cell.com/trends/genetics/abstract/S0168-9525(17)30111-7 (2023) (Sept. 2017).

13. Bates, T. C. et al. Genetic and environmental bases of reading and spelling: A unified genetic dual route model. en. Reading and Writing 20, 147–171. issn: 1573-0905. 10.1007/s11145-006-9022-1 (2023) (Feb. 2007).

14. Doust, C. et al. Discovery of 42 genome-wide significant loci associated with dyslexia. en. Nature Genetics 54. Number: 11 Publisher: Nature Publishing Group, 1621–1629. issn: 1546-1718. https://www.nature.com/articles/s41588-022-01192-y (2023) (Nov. 2022).

15. Williams, L. Z. J. et al. Structural and functional asymmetry of the neonatal cerebral cortex. en. Nature Human Behaviour 7. Number: 6 Publisher: Nature Publishing Group, 942–955. issn: 2397-3374. https://www.nature.com/articles/s41562-023-01542-8 (2023) (June 2023).

16. De Kovel, C. G. F., Carríon-Castillo, A. & Francks, C. A large-scale population study of early life factors influencing left-handedness. en. Scientific Reports 9. Number: 1 Publisher: Nature Publishing Group, 584. issn: 2045-2322. https://www.nature.com/articles/s41598-018-37423-8 (2023) (Jan. 2019).

17. Francks, C. Exploring human brain lateralization with molecular genetics and genomics. en. Annals of the New York Academy of Sciences 1359. eprint: https://onlinelibrary.wiley.com/doi/pdf/10.1111/nyas.12770, 1–13. issn: 1749-6632. https://onlinelibrary.wiley.com/doi/abs/10.1111/nyas.12770 (2023) (2015).

18. Sha, Z. et al. The genetic architecture of structural left–right asymmetry of the human brain. en. Nature Human Behaviour 5. Number: 9 Publisher: Nature Publishing Group, 1226–1239. issn: 2397-3374. https://www.nature.com/articles/s41562-021-01069-w (2023) (Sept. 2021).

19. Sha, Z. et al. Handedness and its genetic influences are associated with structural asymmetries of the cerebral cortex in 31,864 individuals. Proceedings of the National Academy of Sciences 118. Publisher: Proceedings of the National Academy of Sciences, e2113095118. https://www.pnas.org/doi/abs/10.1073/pnas.2113095118 (2023) (Nov. 2021).

20. Wiberg, A. et al. Handedness, language areas and neuropsychiatric diseases: insights from brain imaging and genetics. Brain 142, 2938–2947. issn: 0006-8950. 10.1093/brain/awz257 (2023) (Oct. 2019).

21. Tavor, I. et al. Task-free MRI predicts individual differences in brain activity during task performance. Science 352. Publisher: American Association for the Advancement of Science, 216–220. https://www.science.org/doi/10.1126/science.aad8127 (2023) (Apr. 2016).

22. Joliot, M., Tzourio-Mazoyer, N. & Mazoyer, B. Intra-hemispheric intrinsic connectivity asymmetry and its relationships with handedness and language Lateralization. Neuropsychologia. The Neural Bases of Hemispheric Specialisation 93, 437–447. issn: 0028-3932. https://www.sciencedirect.com/science/article/pii/S0028393216300768 (2023) (Dec. 2016).

23. Labache, L., Ge, T., Yeo, B. T. T. & Holmes, A. J. Language network lateralization is reflected throughout the macroscale functional organization of cortex. eng. Nature Communications 14, 3405. issn: 2041-1723 (June 2023).

24. Smith, S. M. et al. Correspondence of the brain’s functional architecture during activation and rest. Proceedings of the National Academy of Sciences 106. Publisher: Proceedings of the National Academy of Sciences, 13040– 13045. https://www.pnas.org/doi/full/10.1073/pnas.0905267106 (2023) (Aug. 2009).

25. Margulies, D. S. et al. Situating the default-mode network along a principal gradient of macroscale cortical organization. Proceedings of the National Academy of Sciences 113. Publisher: Proceedings of the National Academy of Sciences, 12574–12579. https://www.pnas.org/doi/abs/10.1073/pnas.1608282113 (2023) (Nov. 2016).

26. Bradshaw, A. R., Thompson, P. A., Wilson, A. C., Bishop, D. V. M. & Woodhead, Z. V. J. Measuring language lateralisation with different language tasks: a systematic review. en. PeerJ 5. Publisher: PeerJ Inc., e3929. issn: 2167-8359. https://peerj.com/articles/3929 2023) (Oct. 2017).

27. Elliott, L. T. et al. Genome-wide association studies of brain imaging phenotypes in UK Biobank. en. Nature 562. Number: 7726 Publisher: Nature Publishing Group, 210–216. issn: 1476-4687. https://www.nature.com/articles/s41586-018-0571-7 (2023) (Oct. 2018).

28. Mekki, Y. et al. The genetic architecture of language functional connectivity. eng. NeuroImage 249, 118795. issn: 1095-9572 (Apr. 2022).

29. Yarkoni, T., Poldrack, R. A., Nichols, T. E., Van Essen, D. C. & Wager, T. D. Large-scale automated synthesis of human functional neuroimaging data. en. Nature Methods 8. Number: 8 Publisher: Nature Publishing Group, 665–670. issn: 1548-7105. https://www.nature.com/articles/nmeth.1635 (2023) (Aug. 2011).

30. Joliot, M. et al. AICHA: An atlas of intrinsic connectivity of homotopic areas. en. Journal of Neuroscience Methods 254, 46–59. issn: 01650270. https://linkinghub.elsevier.com/retrieve/pii/S0165027015002678 (2022) (Oct. 2015).

31. Carrion-Castillo, A. et al. Genome sequencing for rightward hemispheric language dominance. en. Genes, Brain and Behavior 18. eprint: https://onlinelibrary.wiley.com/doi/pdf/10.1111/gbb.12572, e12572. issn: 1601-183X. https://onlinelibrary.wiley.com/doi/abs/10.1111/gbb.12572 (2023) (2019).

32. Backman, J. D., et al. Exome sequencing and analysis of 454,787 UK Biobank participants. en. Nature 599. Number: 7886 Publisher: Nature Publishing Group, 628–634. issn: 1476-4687. https://www.nature.com/articles/s41586-021-04103-z (2023) (Nov. 2021).

33. Karczewski, K. J. et al. Systematic single-variant and gene-based association testing of thousands of phenotypes in 394,841 UK Biobank exomes. en. Cell Genomics 2, 100168. issn: 2666979X. https://linkinghub.elsevier.com/retrieve/pii/S2666979X22001100 (2023) (Sept. 2022).

34. Bycroft, C., et al. The UK Biobank resource with deep phenotyping and genomic data. en. Nature 562. Number: 7726 Publisher: Nature Publishing Group, 203–209. issn: 1476-4687. https://www.nature.com/articles/s41586-018-0579-z (2023) (Oct. 2018).

35. Van der Meer, D. et al. Understanding the genetic determinants of the brain with MOSTest. en. Nature Communications 11. Number: 1 Publisher: Nature Publishing Group, 3512. issn: 2041-1723. https://www.nature.com/articles/s41467-020-17368-1 (2023) (July 2020).

36. Watanabe, K., Taskesen, E., van Bochoven, A. & Posthuma, D. Functional mapping and annotation of genetic associations with FUMA. en. Nature Communications 8. Number: 1 Publisher: Nature Publishing Group, 1826. issn: 2041-1723. https://www.nature.com/articles/s41467-017-01261-5 (2023) (Nov. 2017).

37. Leeuw, C. A. d., Mooij, J. M., Heskes, T. & Posthuma, D. MAGMA: Generalized Gene-Set Analysis of GWAS Data. en. PLOS Computational Biology 11. Publisher: Public Library of Science, e1004219. issn: 1553-7358. https://journals.plos.org/ploscompbiol/article?id=10.1371/journal.pcbi.1004219 (2023) (Apr. 2015).

38. Sunkin, S. M. et al. Allen Brain Atlas: an integrated spatio-temporal portal for exploring the central nervous system. Nucleic Acids Research 41, D996–D1008. issn: 0305-1048. 10.1093/nar/gks1042 (2023) (Jan. 2013).

39. Subramanian, A. et al. Gene set enrichment analysis: A knowledge-based approach for interpreting genomewide expression profiles. Proceedings of the National Academy of Sciences 102. Publisher: Proceedings of the National Academy of Sciences, 15545–15550. https://www.pnas.org/doi/10.1073/pnas.0506580102 (2023) (Oct. 2005).

40. Liberzon, A. et al. Molecular signatures database (MSigDB) 3.0. Bioinformatics 27, 1739–1740. issn: 1367-4803. 10.1093/bioinformatics/btr260 (2023) (June 2011).

41. THE GTEX CONSORTIUM. The GTEx Consortium atlas of genetic regulatory effects across human tissues. Science 369. Publisher: American Association for the Advancement of Science, 1318–1330. https://www.science.org/doi/10.1126/science.aaz1776 (2023) (Sept. 2020).

42. Ge, T., Chen, C.-Y., Ni, Y., Feng, Y.-C. A. & Smoller, J. W. Polygenic prediction via Bayesian regression and continuous shrinkage priors. en. Nature Communications 10. Number: 1 Publisher: Nature Publishing Group, 1776. issn: 2041-1723. https://www.nature.com/articles/s41467-019-09718-5 (2023) (Apr. 2019).

43. Mbatchou, J. et al. Computationally efficient whole-genome regression for quantitative and binary traits. en. Nature Genetics 53. Number: 7 Publisher: Nature Publishing Group, 1097–1103. issn: 1546-1718. https://www.nature.com/articles/s41588-021-00870-7 (2023) (July 2021).

44. Lee, S. et al. Optimal Unified Approach for Rare-Variant Association Testing with Application to Small-Sample Case-Control Whole-Exome Sequencing Studies. en. The American Journal of Human Genetics 91, 224–237. issn: 00029297. https://linkinghub.elsevier.com/retrieve/pii/S0002929712003163 (2023) (Aug. 2012).

45. Szustakowski, J. D. et al. Advancing human genetics research and drug discovery through exome sequencing of the UK Biobank. en. Nature Genetics 53. Number: 7 Publisher: Nature Publishing Group, 942–948. issn: 1546-1718. https://www.nature.com/articles/s41588-021-00885-0 (2023) (July 2021).

46. Pasquale, E. B. Eph-Ephrin Bidirectional Signaling in Physiology and Disease. en. Cell 133, 38–52. issn: 0092-8674. https://www.sciencedirect.com/science/article/pii/S0092867408003863 (2023) (Apr. 2008).

47. Gibson, D. A. & Ma, L. Developmental regulation of axon branching in the vertebrate nervous system. eng. Development (Cambridge, England) 138, 183–195. issn: 1477-9129 (Jan. 2011).

48. Gerstmann, K. & Zimmer, G. The role of the Eph/ephrin family during cortical development and cerebral malformations. en-US. Medical Research Archives 6. issn: 2375-1924. https://esmed.org/MRA/mra/article/view/1694 (2023) (Mar. 2018).

49. Zhao, B. et al. Common variants contribute to intrinsic human brain functional networks. en. Nature Genetics 54. Number: 4 Publisher: Nature Publishing Group, 508–517. issn: 1546-1718. https://www.nature.com/articles/s41588-022-01039-6 (2023) (Apr. 2022).

50. Sha, Z., Schijven, D., Fisher, S. E. & Francks, C. Genetic architecture of the white matter connectome of the human brain. eng. Science Advances 9, eadd2870. issn: 2375-2548 (Feb. 2023).

51. Fagerberg, L. et al. Analysis of the human tissue-specific expression by genome-wide integration of transcriptomics and antibody-based proteomics. eng. Molecular & cellular proteomics: MCP 13, 397–406. issn: 1535-9484 (Feb. 2014).

52. Fan, C. C. et al. Multivariate genome-wide association study on tissue-sensitive diffusion metrics highlights pathways that shape the human brain. en. Nature Communications 13. Number: 1 Publisher: Nature Publishing Group, 2423. issn: 2041-1723. https://www.nature.com/articles/s41467-022-30110-3 (2023) (May 2022).

53. Nalls, M. A. et al. Identification of novel risk loci, causal insights, and heritable risk for Parkinson’s disease: a meta-analysis of genome-wide association studies. eng. The Lancet. Neurology 18, 1091–1102. issn: 1474-4465 (Dec. 2019).

54. Trubetskoy, V. et al. Mapping genomic loci implicates genes and synaptic biology in schizophrenia. eng. Nature 604, 502–508. issn: 1476-4687 (Apr. 2022).

55. Sollis, E. et al. The NHGRI-EBI GWAS Catalog: knowledgebase and deposition resource. Nucleic Acids Research 51, D977–D985. issn: 0305-1048. 10.1093/nar/gkac1010 (2023) (Jan. 2023).

56. Cuellar-Partida, G. et al. Genome-wide association study identifies 48 common genetic variants associated with handedness. en. Nature Human Behaviour 5. Number: 1 Publisher: Nature Publishing Group, 59–70. issn: 2397-3374. https://www.nature.com/articles/s41562-020-00956-y (2023) (Jan. 2021).

57. Knecht, S. et al. Behavioural relevance of atypical language lateralization in healthy subjects. Brain 124, 1657– 1665. issn: 0006-8950. 10.1093/brain/124.8.1657 (2023) (Aug. 2001).

58. Bruckert, L. Is language laterality related to language abilities? English. http://purl.org/dc/dcmitype/Text (University of Oxford, 2016). https://ora.ox.ac.uk/objects/uuid:05e80d0d-8d0b-4cb2-8f94-22763603fab5 (2023).

59. Mellet, E. et al. Weak language lateralization affects both verbal and spatial skills: An fMRI study in 297 subjects. Neuropsychologia 65, 56–62. issn: 0028-3932. https://www.sciencedirect.com/science/article/pii/S0028393214003704 (2023) (Dec. 2014).

60. Leonard, C. M. & Eckert, M. A. Asymmetry and Dyslexia. Developmental Neuropsychology 33. Publisher: Routledge eprint: 10.1080/87565640802418597, 663–681. issn: 8756-5641. 10.1080/87565640802418597 (2023) (Nov. 2008).

61. Richlan, F., Kronbichler, M. & Wimmer, H. Meta-analyzing brain dysfunctions in dyslexic children and adults. NeuroImage 56, 1735–1742. issn: 1053-8119. https://www.sciencedirect.com/science/article/pii/S1053811911001960 (2023) (June 2011).

62. Van der Mark, S. et al. The left occipitotemporal system in reading: Disruption of focal fMRI connectivity to left inferior frontal and inferior parietal language areas in children with dyslexia. NeuroImage 54, 2426–2436. issn: 1053-8119. https://www.sciencedirect.com/science/article/pii/S1053811910012930 (2023) (Feb. 2011).

63. Kershner, J. R. Neuroscience and education: Cerebral lateralization of networks and oscillations in dyslexia. Laterality 25. Publisher: Routledge eprint: 10.1080/1357650X.2019.1606820, 109–125. issn: 1357-650X. 10.1080/1357650X.2019.1606820 (2023) (Jan. 2020).

64. Zago, L. et al. Predicting hemispheric dominance for language production in healthy individuals using support vector machine. en. Human Brain Mapping 38. eprint: https://onlinelibrary.wiley.com/doi/pdf/10.1002/hbm.23770, 5871–5889. issn: 1097-0193. https://onlinelibrary.wiley.com/doi/abs/10.1002/hbm.23770 (2023) (2017).

65. Sha, Z., Schijven, D. & Francks, C. Patterns of brain asymmetry associated with polygenic risks for autism and schizophrenia implicate language and executive functions but not brain masculinization. en. Molecular Psychiatry 26. Number: 12 Publisher: Nature Publishing Group, 7652–7660. issn: 1476-5578. https://www.nature.com/articles/s41380-021-01204-z (2023) (Dec. 2021).

66. Wang, H.-T. et al. Finding the needle in a high-dimensional haystack: Canonical correlation analysis for neuroscientists. en. NeuroImage 216, 116745. issn: 1053-8119. https://www.sciencedirect.com/science/article/pii/S1053811920302329 (2023) (Aug. 2020).

67. Smith, S. M. et al. A positive-negative mode of population covariation links brain connectivity, demographics and behavior. en. Nature Neuroscience 18. Number: 11 Publisher: Nature Publishing Group, 1565–1567. issn: 1546-1726. https://www.nature.com/articles/nn.4125 (2023) (Nov. 2015).

68. Lee, J. J. et al. Gene discovery and polygenic prediction from a genome-wide association study of educational attainment in 1.1 million individuals. eng. Nature Genetics 50, 1112–1121. issn: 1546-1718 (July 2018).

69. Gaudet, P., Livstone, M. S., Lewis, S. E. & Thomas, P. D. Phylogenetic-based propagation of functional annotations within the Gene Ontology consortium. eng. Briefings in Bioinformatics 12, 449–462. issn: 1477-4054 (Sept. 2011).

70. Bijsterbosch, J. et al. Challenges and future directions for representations of functional brain organization. en. Nature Neuroscience 23. Number: 12 Publisher: Nature Publishing Group, 1484–1495. issn: 1546-1726. https://www.nature.com/articles/s41593-020-00726-z (2023) (Dec. 2020).

71. Bijsterbosch, J. D., Valk, S. L., Wang, D. & Glasser, M. F. Recent developments in representations of the connectome. en. NeuroImage 243, 118533. issn: 1053-8119. https://www.sciencedirect.com/science/article/pii/S1053811921008065 (2023) (Nov. 2021).

72. Bignardi, G. et al. Pervasive inter-individual differences in the sensorimotor-association axis of cortical organization en. preprint (Neuroscience, July 2023). http://biorxiv.org/lookup/doi/10.1101/2023.07.13.548817 (2023).

73. Llera, A., Wolfers, T., Mulders, P. & Beckmann, C. F. Inter-individual differences in human brain structure and morphology link to variation in demographics and behavior. en. eLife 8, e44443. issn: 2050-084×. https://elifesciences.org/articles/44443 (2023) (July 2019).

74. Pang, J. C., et al. Geometric constraints on human brain function. en. Nature 618. Number: 7965 Publisher: Nature Publishing Group, 566–574. issn: 1476-4687. https://www.nature.com/articles/s41586-023-06098-1 (2023) (June 2023).

75. Suárez, L. E., Markello, R. D., Betzel, R. F. & Misic, B. Linking Structure and Function in Macroscale Brain Networks. English. Trends in Cognitive Sciences 24. Publisher: Elsevier, 302–315. issn: 1364-6613, 1879-307X. https://www.cell.com/trends/cognitive-sciences/abstract/S1364-6613(20)30026-7 (2023) (Apr. 2020).

76. Huffman, J. E. Examining the current standards for genetic discovery and replication in the era of mega-biobanks. en. Nature Communications 9. Number: 1 Publisher: Nature Publishing Group, 5054. issn: 2041-1723. https://www.nature.com/articles/s41467-018-07348-x (2023) (Nov. 2018).

77. Fry, A. et al. Comparison of Sociodemographic and Health-Related Characteristics of UK Biobank Participants With Those of the General Population. American Journal of Epidemiology 186, 1026–1034. issn: 0002-9262. 10.1093/aje/kwx246 (2023) (Nov. 2017).

78. Sudlow, C. et al. UK Biobank: An Open Access Resource for Identifying the Causes of a Wide Range of Complex Diseases of Middle and Old Age. en. PLOS Medicine 12. Publisher: Public Library of Science, e1001779. issn: 1549-1676. https://journals.plos.org/plosmedicine/article?id=10.1371/journal.pmed.1001779 (2023) (Mar. 2015).

79. Alfaro-Almagro, F. et al. Image processing and Quality Control for the first 10,000 brain imaging datasets from UK Biobank. en. NeuroImage 166, 400–424. issn: 1053-8119. https://www.sciencedirect.com/science/article/pii/S1053811917308613 (2023) (Feb. 2018).

80. Miller, K. L. et al. Multimodal population brain imaging in the UK Biobank prospective epidemiological study. en. Nature Neuroscience 19. Number: 11 Publisher: Nature Publishing Group, 1523–1536. issn: 1546-1726. https://www.nature.com/articles/nn.4393 (2023) (Nov. 2016).

81. Van Hout, C. V. et al. Exome sequencing and characterization of 49,960 individuals in the UK Biobank. en. Nature 586. Number: 7831 Publisher: Nature Publishing Group, 749–756. issn: 1476-4687. https://www.nature.com/articles/s41586-020-2853-0 (2023) (Oct. 2020).

82. Jenkinson, M., Bannister, P., Brady, M. & Smith, S. Improved Optimization for the Robust and Accurate Linear Registration and Motion Correction of Brain Images. en. NeuroImage 17, 825–841. issn: 1053-8119. https://www.sciencedirect.com/science/article/pii/S1053811902911328 (2023) (Oct. 2002).

83. Beckmann, C. & Smith, S. Probabilistic independent component analysis for functional magnetic resonance imaging. IEEE Transactions on Medical Imaging 23. Conference Name: IEEE Transactions on Medical Imaging, 137–152. issn: 1558-254× (Feb. 2004).

84. Griffanti, L. et al. ICA-based artefact removal and accelerated fMRI acquisition for improved resting state network imaging. en. NeuroImage 95, 232–247. issn: 1053-8119. https://www.sciencedirect.com/science/article/pii/S1053811914001815 (2023) (July 2014).

85. Salimi-Khorshidi, G. et al. Automatic denoising of functional MRI data: Combining independent component analysis and hierarchical fusion of classifiers. en. NeuroImage 90, 449–468. issn: 1053-8119. https://www.sciencedirect.com/science/article/pii/S1053811913011956 (2023) (Apr. 2014).

86. Jenkinson, M., Beckmann, C. F., Behrens, T. E. J., Woolrich, M. W. & Smith, S. M. FSL. en. NeuroImage. 20 YEARS OF fMRI 62, 782–790. issn: 1053-8119. https://www.sciencedirect.com/science/article/pii/S1053811911010603 (2023) (Aug. 2012).

87. Fischl, B. FreeSurfer. en. NeuroImage. 20 YEARS OF fMRI 62, 774–781. issn: 1053-8119. https://www.sciencedirect.com/science/article/pii/S1053811912000389 (2023) (Aug. 2012).

88. Yang, J., Lee, S. H., Goddard, M. E. & Visscher, P. M. GCTA: A Tool for Genome-wide Complex Trait Analysis. en. The American Journal of Human Genetics 88, 76–82. issn: 0002-9297. https://www.sciencedirect.com/science/article/pii/S0002929710005987 (2023) (Jan. 2011).

89. Ni, G. et al. A Comparison of Ten Polygenic Score Methods for Psychiatric Disorders Applied Across Multiple Cohorts. eng. Biological Psychiatry 90, 611–620. issn: 1873-2402 (Nov. 2021).

90. Zheutlin, A. B. et al. Penetrance and Pleiotropy of Polygenic Risk Scores for Schizophrenia in 106,160 Patients Across Four Health Care Systems. eng. The American Journal of Psychiatry 176, 846–855. issn: 1535-7228 (Oct. 2019).

91. Cingolani, P. et al. A program for annotating and predicting the effects of single nucleotide polymorphisms, SnpEff: SNPs in the genome of Drosophila melanogaster strain w1118; iso-2; iso-3. eng. Fly 6, 80–92. issn: 1933-6942 (2012).

92. Liu, X., Li, C., Mou, C., Dong, Y. & Tu, Y. dbNSFP v4: a comprehensive database of transcript-specific functional predictions and annotations for human nonsynonymous and splice-site SNVs. eng. Genome Medicine 12, 103. issn: 1756-994× (Dec. 2020).

93. Rentzsch, P., Witten, D., Cooper, G. M., Shendure, J. & Kircher, M. CADD: predicting the deleteriousness of variants throughout the human genome. Nucleic Acids Research 47, D886–D894. issn: 0305-1048. 10.1093/nar/gky1016 (2023) (Jan. 2019).

